# Gene regulatory innovations from transposable elements in mammalian cerebellum development

**DOI:** 10.1101/2025.10.15.682527

**Authors:** Tetsuya Yamada, Mari Sepp, Ioannis Sarropoulos, Henrik Kaessmann

## Abstract

Transposable elements (TEs) are hypothesized to have driven gene regulatory innovation, yet their contributions to primate brain development at the cell type level remain largely unexplored. Using single-cell multiomics data for the cerebellum from human, macaque, marmoset, and mouse, we revealed that TE contributions to different cell types are shaped by differences in evolutionary constraint across cell types, as well as the preferential co-option of certain TEs in specific cell states. Using a sequence-based deep-learning model predicting cell-type-specific chromatin accessibility, we systematically assessed the co-option potential of TEs into cerebellar gene regulatory networks, identifying seventeen TE fragments with complex regulatory sequences in their ancestral states that facilitate their co-option as cell-type-specific cis-regulatory elements. Primate-specific HERVL (human endogenous retrovirus L) is predominantly accessible in differentiating granule cells and other rhombic-lip-derived neuroblasts during hindbrain development, and contributes to species-specific gene expression patterns. Our study uncovers a pivotal role for TEs in the evolution of the cerebellum. More generally, it demonstrates how TEs can be directly co-opted into the gene regulatory networks of specific cell types and introduces a generalizable analytical framework for dissecting their contribution to mammalian regulatory evolution.

## INTRODUCTION

The genetic basis of brain evolution, particularly in the primate lineage, remains to be fully elucidated. The cerebellum has undergone significant expansion in neuron numbers alongside the neocortex during primate evolution, with cerebellar granule cells—which develop from the upper rhombic lip in the hindbrain and constitute approximately 80% of all neurons in the human brain—playing a central role in this expansion^1,2^. Gene regulatory changes affecting gene expression are considered central to such evolutionary innovations^3^, as they can modify specific cellular processes without disrupting essential functions^4^. These regulatory changes occur predominantly through modifications to cis-regulatory elements (CREs)^5^, such as promoters and enhancers, which are bound by transcription factors (TFs) and regulate genes in a cell-type-specific manner^6^. Many evolutionary changes arise from single-nucleotide substitutions and small insertions or deletions (indels).

Transposable elements (TEs) offer another mechanism for regulatory innovation^7–9^. These DNA sequences, capable of self-replication and genomic translocation, have long been implicated in transcriptional regulation by rewiring gene regulatory networks through the expansion of copies harboring TF binding sites. This model of TE-driven regulatory evolution was already postulated by Barbara McClintock in the 1950s, who first identified TEs in maize and referred to them as ‘controlling elements’^10^, a concept that was later expanded as the ‘gene-battery’ model by Britten and Davidson^11–13^. Indeed, recent genome-wide studies on CREs across different tissues have revealed that TEs overlap approximately one-quarter of CREs^14–16^. Furthermore, some TEs are enriched for regulatory sequences, likely due to pre-existing TF binding sites in their ancestral sequences^14,17–20^. These lines of evidence suggest an important role of TEs in gene regulatory innovation.

However, the extent and mechanisms of TE contributions to regulatory innovations remain debated. First, despite comprising 45% to 70% of the human genome^21,22^, TEs appear to be underrepresented in CREs compared to their overall genomic abundance^23^. Second, the observed association between TEs and regulatory regions may partly reflect their inherent insertion site preferences, given that TEs from certain families tend to insert into open chromatin regions^24–26^. Third, TE insertions that affect gene expression often face rapid elimination through purifying selection, suggesting that their large-scale genomic alterations may be too disruptive to be readily co-opted for regulatory innovation^27^. Finally, most previous studies lacked cellular resolution, limiting our understanding of TE contributions to gene regulation, which fundamentally operates at the level of individual cell types. This limitation is particularly relevant for tissues with high cellular complexity, such as the developing brain.

Our previous work characterized CRE and gene expression innovations across different cell types in mammalian cerebellar development^28–30^. We developed a deep-learning model, DeepCeREvo, which learned the sequence grammar of cerebellar cell-type-specific CREs and applied it across the genomes of 240 mammalian species to understand the genetic basis of CRE and gene expression evolution in the lineage leading to humans^30^. This analysis identified hundreds of CRE innovations in each cell type, which occurred at different times points during evolution and include human-specific events, and characterized single nucleotide substitutions and small indels potentially underlying these innovations. However, the contributions of TEs to these CRE innovations and the overall regulatory landscape in cerebellar cell types remains unexplored.

In this study, we combined single-cell multi-omics datasets of developing cerebellar cell types from human, macaque, marmoset, and mouse with our sequence-based deep-learning model to systematically investigate how TEs shape gene regulatory evolution in cerebellar development.

## RESULTS

### TE contributions to species- and cell-type-specific cis-regulatory elements in mammalian cerebellum development

To assess the contributions of transposable elements (TEs) to the cis-regulatory landscape in mammalian cerebellum development, we intersected 557,491 candidate cis-regulatory elements (cCREs) accessible in the human cerebellum^30^ with annotated TEs^31^. Overall, 18.1% of human cCREs overlap with TEs. This overlap is lower than expected based on the abundance of TEs in the human genome (45%)^21^, implying a depletion of TEs amongst cCREs^16,23^. While all classes of cCREs show significant depletion of TE overlap compared to genomic background, the extent of this depletion varies, with distal elements exhibiting the highest TE overlap (23.8%) and exonic elements the lowest (5.3%), followed by promoters (Fig. 1a), consistent with previous studies^15,23^. This pattern supports the notion that TE insertions in functionally critical and evolutionarily constrained regions primarily have deleterious effects^32^.

**Fig. 1:**
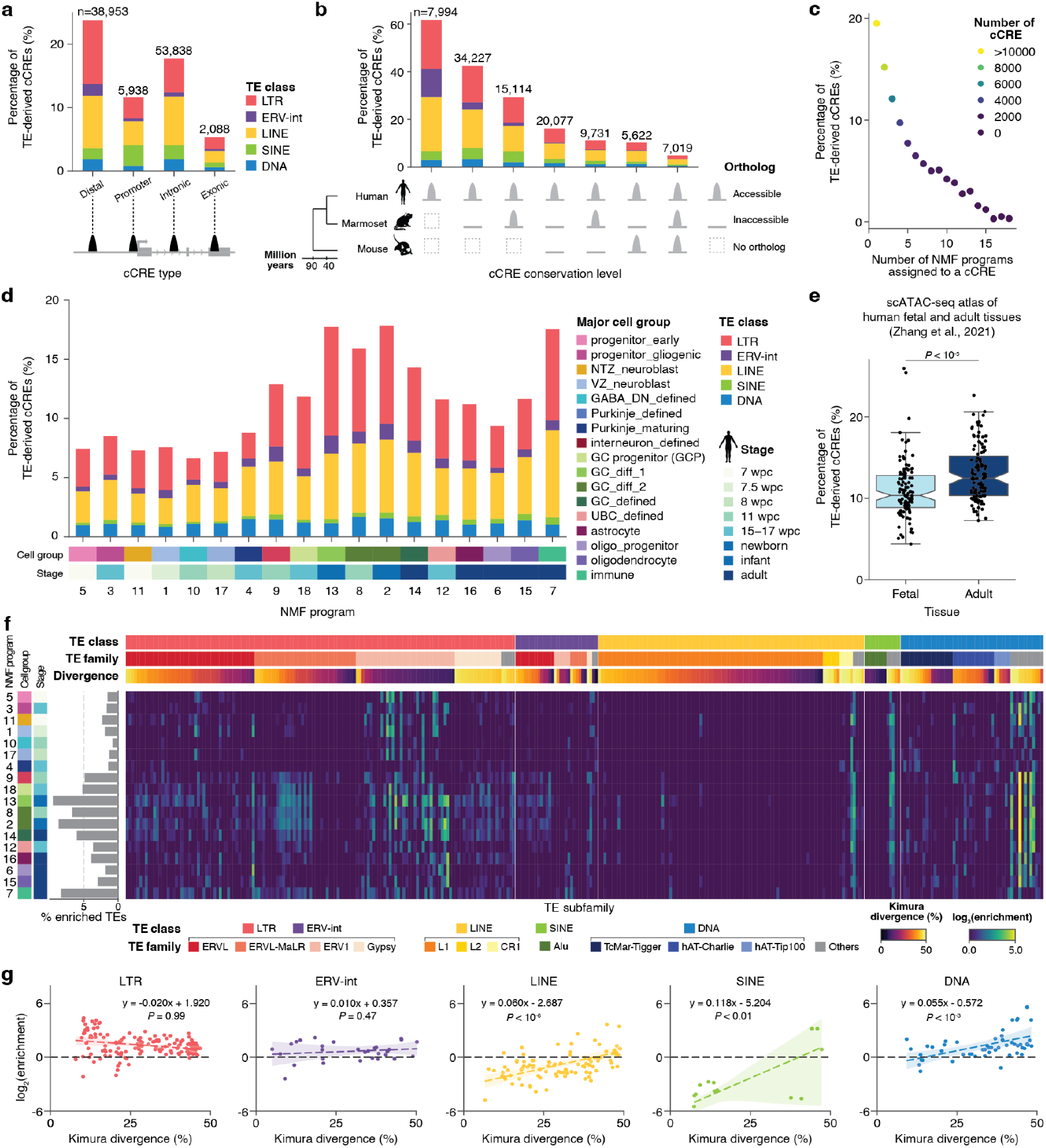
Contributions of transposable elements to cCREs in human cerebellar development. **a, b**, Percentage of TE-derived cCREs in human cerebellar development across genomic contexts (**a**) or sequence and accessibility conservation between human, marmoset, and mouse (**b**). **c**, Relationship between cCRE accessibility specificity across NMF programs and percentage of TE-derived cCREs. **d**, Percentage of NMF program-specific cCREs that originate from TEs across human cerebellar cell types and developmental stages. **e**, Percentage of TE-derived cCREs in cell types from fetal or adult tissues across the human body, with each point representing a cell type. Data from Zhang et al., 2021^41^ reanalyzed. **f**, Subfamily-specific TE enrichment within NMF program-specific cCREs. (Left) bar plot indicates the proportion of each enriched TE subfamily contributing to TE-derived cCREs. **g**, Correlation between evolutionary age (median Kimura divergence) and TE enrichment in NMF program-specific cCREs (maximum log_2_ enrichment across NMF programs) for each TE subfamily, stratified by TE class. Linear regression equations and *P*-values are shown with 95% confidence intervals. DN, deep nuclei neuron; ERV-int, internal sequence of endogenous retrovirus; GC, granule cell; LTR, long terminal repeat of endogenous retrovirus; NMF, non-negative matrix factorization; NTZ, nuclear transitory zone; UBC, unipolar brush cell; VZ, ventricular zone; wpc, weeks post conception.

Despite the overall depletion, TE-associated cCREs are disproportionately represented among species- or lineage-specific regulatory elements. TEs contribute to 62% of human-specific and 29% of primate-specific cCREs, showing significant overrepresentation compared to all human cCREs (Fig. 1b; Fisher’s exact *P* < 10^−30^). These contributions of TEs to primate-specific cCREs are lower than previous estimations of approximately 80%^33,34^, likely due to the more conservative TE-cCRE overlap criteria (more than 50% of cCRE length) used in this study. On the other hand, cCREs conserved across mammals are significantly underrepresented in TE overlaps (5%) compared to all human cCREs, though the contributions of older TEs to conserved cCREs may be underestimated due to the challenges in annotating highly degenerated TEs^35^.

To determine whether this pattern generalizes across mammalian species, we intersected cerebellar cCREs from marmoset (n = 505,724) and mouse (n = 498,119) with TE annotations from their respective genomes. The results confirmed a consistent overrepresentation of TEs in species-specific elements: 63% of marmoset-specific cCREs and 34% of mouse-specific cCREs derive from TEs, whereas less than 5% of cCREs with conserved accessibility across species are associated with TEs in both marmoset and mouse (Extended Data Fig. 1a,b). These findings confirm that TE-derived cCREs are enriched in more recently emerged regulatory elements across mammals^36–39^.

To characterize cell-type-specific patterns of TE integration into cCREs, we analyzed cCREs associated with 18 distinct programs derived from non-negative matrix factorization (NMF)^30^. These programs, computed by decomposing chromatin accessibility matrices across cell types and developmental stages, capture the major patterns of regulatory activity characteristic of particular cell populations at defined developmental timepoints, thereby enabling systematic analysis of cell-type-specific TE contributions throughout cerebellar development. TE-derived sequences constitute 20% of intergenic cCREs specific to a single NMF program but, on average, only 2.6% of those assigned to more than half of the 18 NMF programs, demonstrating significant depletion of TEs in broadly accessible cCREs relative to cell-state-specific elements (Fig. 1c), consistent with previous multi-tissue epigenomic studies^15,16,23,40^. This trend may reflect the fact that regulatory elements active in many cell types face stronger evolutionary constraints and require more complex regulatory sequences to coordinate function across diverse cellular contexts, making such elements more challenging to evolve through TE co-option.

The contributions of TEs to cell-type-specific cCREs varies markedly across different cell types in cerebellar development. We observed relative overrepresentation of TEs in cCREs specific to differentiating and mature granule cells (NMF programs 13, 8, 2, and 14) and microglia (NMF program 7) (Fisher’s exact *P* < 10^−28^), whereas cCREs associated with earlier stages of cerebellar development (NMF programs 5, 3, 11, 1, 10, 17, and 4) show significant depletion of TEs (Fig. 1d; Fisher’s exact *P* < 10^−31^). To determine if this pattern applies more broadly across cell types and organs, we re-analyzed data from previous work^41^. The analysis revealed that cCREs in cell types from fetal tissues indeed exhibit significant depletion of TE-derived sequences compared to those from adult tissues (Fig. 1e; *P* < 10^−5^, Mann-Whitney *U* test). Within fetal tissues, we observed tissue-specific variation, with relative underrepresentation of TE-derived cCREs in neural cell types and overrepresentation in placental cell types, consistent with the important role of TEs in placental functions^42,43^ (Extended Data Fig. 2). Together, our results suggest that the differential abundance of TEs across cCREs reflects the varying degrees of evolutionary constraint experienced by different cell types, with greater TE co-option occurring in more specialized, later-developing cell types where regulatory innovation was facilitated by reduced selective constraints^28,44^.

TEs are classified hierarchically into classes, families, and subfamilies based on replication mechanisms and sequence features^31,45^ (Methods). The varied enrichment of TEs in cCREs across cell types may also be explained by the different co-option tendencies of specific TE subfamilies into cell-type-specific regulatory elements^46,47^. To quantify these subfamily-specific patterns, we analyzed the overlaps of TE subfamilies in cCREs across each NMF program and compared their occurrence to baseline frequency in the human genome. TE subfamily ages were inferred by the median Kimura divergence of copies to the corresponding consensus sequence^48^. Our analysis revealed that TE-derived cCREs largely originate from subfamilies without significant enrichment relative to their genomic abundance, with only a small fraction coming from overrepresented subfamilies, ranging from 0.8% (NMF program 10) to 9.5% (NMF program 13) and correlating with the overall TE enrichment across programs (Fig. 1e; Supplementary Table 1). We also observed distinct enrichment patterns across TE subfamilies of different ages and classes. Younger TE subfamilies with lower divergence are more depleted from cCREs than older ones, with the notable exception of long terminal repeat (LTR) and internal sequence of endogenous retroviruses (ERV-int) classes (Fig. 1e,f), as previously reported^14,37,39^. These TEs are preferentially co-opted in the granule cell lineage (NMF programs 13, 8, 2, and 14) and microglia (NMF programs 7), consistent with the overall TE enrichment in these cell types, as well as in progenitor and neuroblast lineages (NMF programs 5, 11, and 1), despite these early developmental cells generally showing significant depletion of TE-derived cCREs (Fig. 1e,f). Collectively, our results suggest that TE contributions to cCREs in different cell types are shaped by both differences in evolutionary constraints across cell types and preferential co-option of specific TE subfamilies.

### Complex cis-regulatory sequences in ancestral TE sequences facilitate co-option of TEs

The preferential co-option of certain TE subfamilies into cell-type-specific cCREs could be driven by pre-existing sequence features capable of functioning as regulatory elements, such as TF binding sites, as demonstrated in several TE subfamilies in specific tissues using reporter assays^17,19,20^. To systematically assess the cis-regulatory potential of ancestral TE sequences in cerebellar cell types, we applied DeepCeREvo, a deep-learning model trained on human and mouse cCRE sequences that predicts NMF program-specific cCREs based on 500 bp nucleotide sequences^30^ (Fig. 2a). For each TE subfamily, we generated 500 bp sliding windows with 100 bp steps across the consensus sequence and analyzed 127 subfamilies that showed at least 10 overlaps with highly variable cCREs in at least one cell type, prioritizing those with high contributions to cell-type-specific chromatin accessibility. We computed cell-type-specific regulatory potential for each fragment using DeepCeREvo and compared it against dinucleotide-shuffled control sequences to determine statistical significance (Fig. 2a; Methods). Additionally, to quantify preferential co-option of these TE fragments, we compared their overlap enrichment with NMF program-specific cCREs versus background cCREs from the whole human body at both fetal and adult stages^41^ (Fig. 2b).

**Fig. 2:**
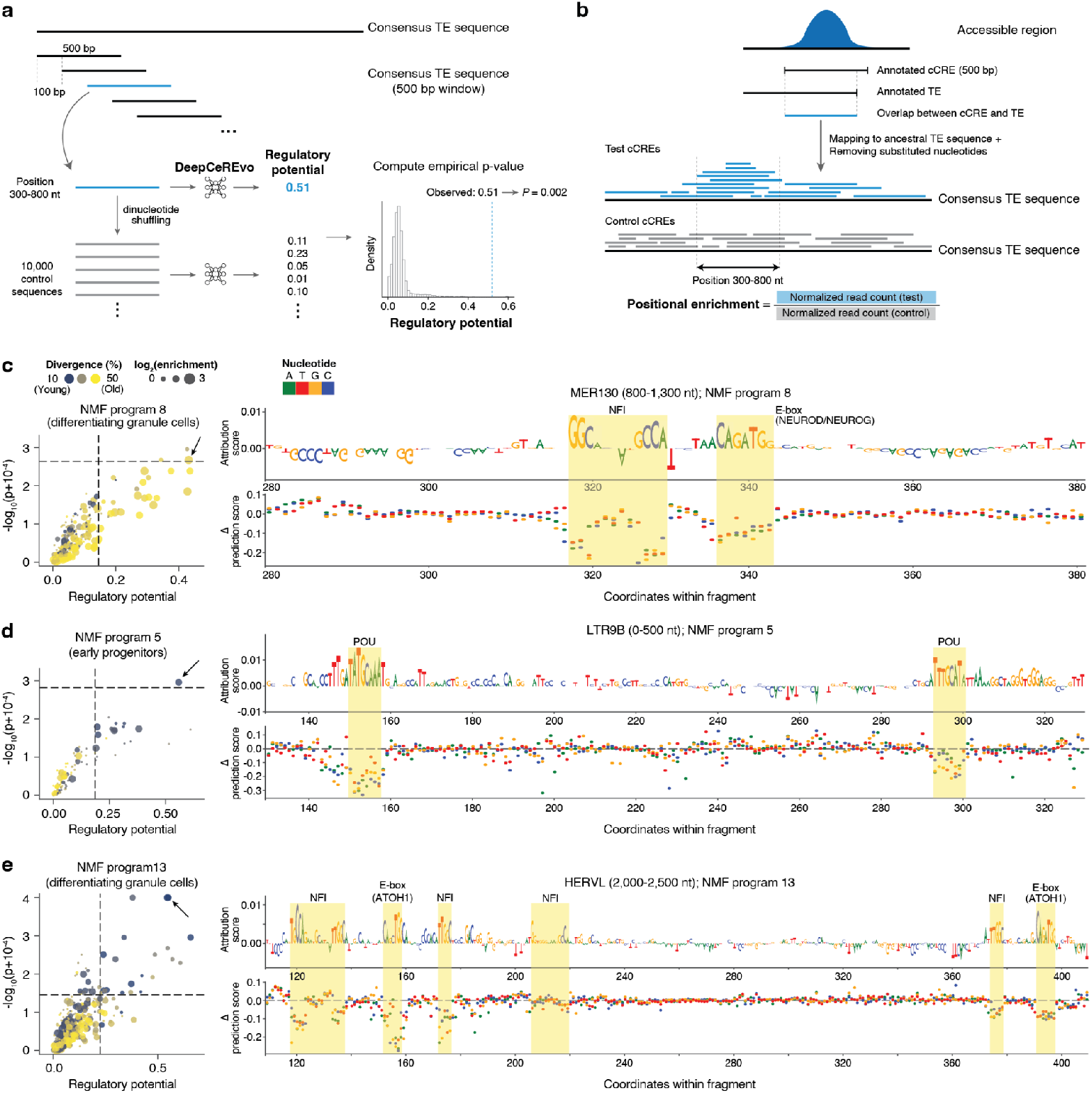
*In silico* screening for transposable elements with high regulatory potential. **a, b**, Schematic overview of the workflows for evaluating the regulatory potential of ancestral (consensus) TE fragments using DeepCeREvo (a) and for assessing enrichment of cell-type-specific cCREs within specific regions of TE fragments compared to background cCRE sets (b). The workflows were applied separately for each NMF/cell state. **c-e**, Identification of TE subfamily fragments with high regulatory potential and high contribution to cell-type-specific cCREs. Left: Screening of TE fragments based on prediction scores and associated *P*-values, alongside enrichment in cell-type-specific cCREs for NMF programs 8, 5, and 13, respectively. The threshold for regulatory potential was set at the 95th percentile. The *P*-value threshold was determined using the Benjamini–Hochberg procedure to control the false discovery rate (FDR) at 5%, adjusted for the effective number of tests. Right: DeepExplainer attribution and *in silico* mutagenesis profiles for top candidates (indicated by arrows in left panels).

Our screening analysis identified 17 non-overlapping TE fragments with high cis-regulatory potential for specific cell types in cerebellum development and enriched overlap with cCREs at corresponding positions in extant copies (Extended Data Fig. 3a; Supplementary Table 2). These include MER130 in NMF program 8 (differentiating granule cells), previously identified as a DNA transposon co-opted into developmental enhancers in the mouse neocortex^49^ (Fig. 2c). Attribution analysis of DeepCeREvo revealed that the reconstructed ancestral sequence of this TE fragment contains an E-box (CAGATGG; NEUROD/NEUROG) and a NFI binding motif (GCCA), both known to be important for brain development, including the differentiation of cerebellar granule cells^28,49–51^ (Fig. 2c).

Our screening also identified multiple fragments from younger, previously uncharacterized TEs, all belonging to the LTR or ERV-int classes, with TF binding sites in their ancestral state that correspond to their enrichment in specific cell type cCREs (Extended Data Fig. 3a). For instance, LTR9B in NMF program 5 (early progenitors) and MER72 in NMF program 1 (ventricular zone neuroblasts) contain POU motifs (ATAT(T/G)CA) (Fig. 2d; Extended Data Fig. 3b). These TFs are known to be active during early embryonic development and their binding motifs are enriched in LTR sequences, facilitating transcription of proviral sequences in germ lines for intergenerational inheritance^47,52,53^ (Fig. 2d). In addition to the TEs containing binding sites for general embryonic regulators, we identified another group with sequences recognised by cell-type-specific TFs. HERVL (human endogenous retrovirus L) shows high regulatory potential in NMF program 13 (differentiating granule cells) with multiple E-box (CAG(C/A)TG; ATOH1) and NFI (GCCA) binding motifs (Fig. 2e). ATOH1 is a TF regulating genes essential for the development of cerebellar granule cells^54–56^. Similarly, ERVL-B4 with multiple NFI motifs is enriched in NMF program 2 (differentiating and defined granule cells) (Extended Data Fig. 3c). Collectively, using a deep learning-based approach, we identified TE fragments with complex cis-regulatory sequences in their ancestral states that facilitate their preferential co-option as CREs in specific cell types.

### HERVL elements are co-opted as regulatory elements in rhombic-lip-derived differentiating neurons

Among the TE subfamilies that exhibited both high regulatory potential and cell-type-specific preferential co-option in our analysis, HERVL stands out as the youngest element, which underwent a major burst of transposition in the primate lineage^57^. To assess HERVL’s specific contribution to the chromatin accessibility landscape in human cerebellar development, we projected the accessibility profiles of all 2,006 annotated HERVL copies in each cell type onto the corresponding consensus sequences. This analysis revealed that HERVL copies are predominantly accessible in differentiating granule cells (GC_diff_1) and developmentally related cell groups, including granule cell progenitors (GCP) and differentiating unipolar brush cells (UBC_diff), with accessibility concentrated in a specific region (position 2,000-2,500 nt) of the HERVL consensus sequence, corresponding to the beginning of the polymerase gene region of the endogenous retrovirus (Fig. 3a). Of the 2,006 annotated HERVL copies in the human genome, 42 (2.1%) were accessible in GC_diff_1 (Fig. 3b). Among these, 20 were also accessible in GCP and 7 in UBC_diff, whereas all other cell groups showed no more than 4 accessible copies.

**Fig. 3:**
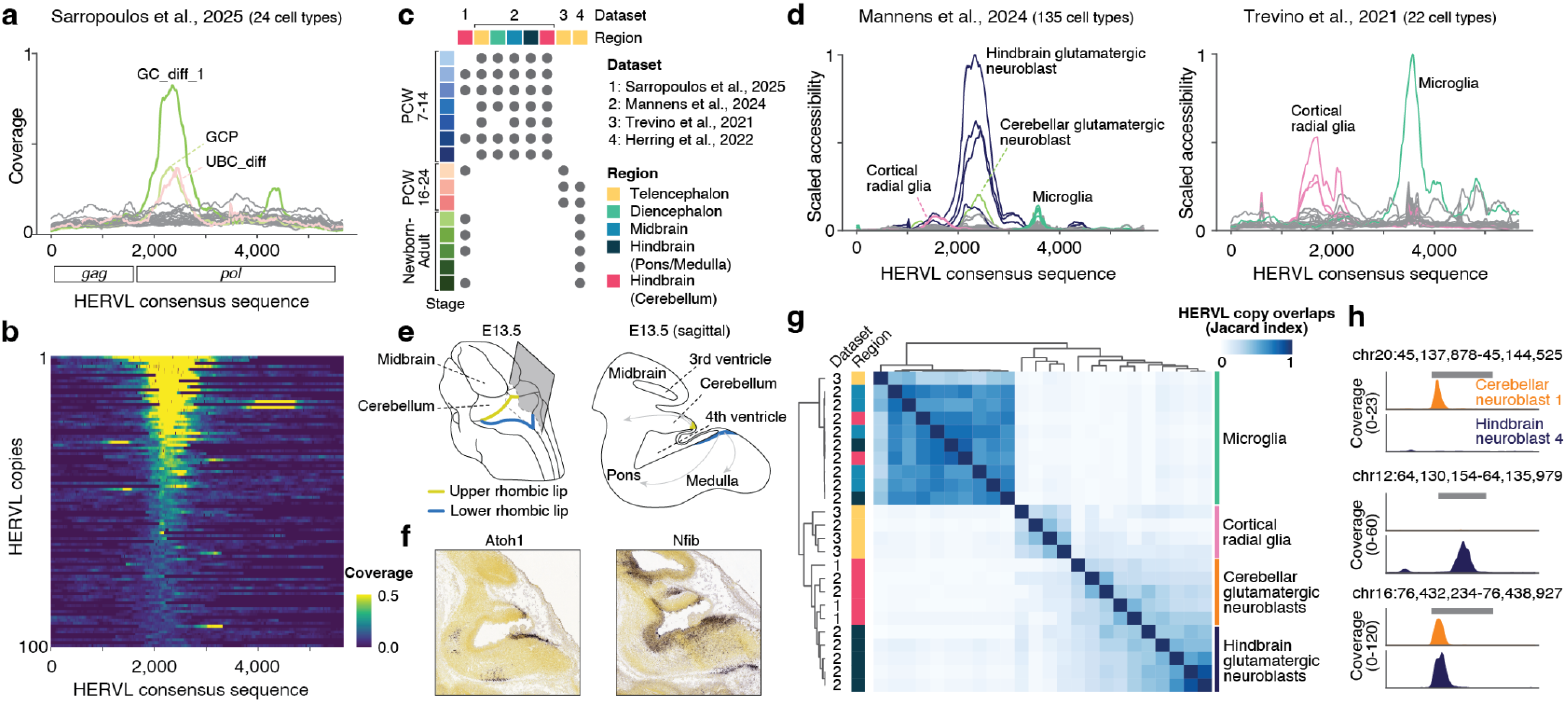
HERVL co-option in CREs specific to rhombic-lip-derived neuroblasts. **a**, Mean chromatin accessibility profiles of the 30 most accessible HERVL copies across 24 cell groups in human cerebellar development, aligned to the HERVL consensus sequence. Regions homologous to gag (group antigen) and pol (polymerase) genes of the HERVL provirus are indicated below the coordinates. **b**, Chromatin accessibility patterns of the 100 most accessible HERVL copies in the 2,000-2,500 nt position in differentiating granule cells (GC_diff_1), aligned to the consensus sequence. **c**, Overview of chromatin accessibility datasets spanning multiple brain regions across developmental stages. **d**, Mean accessibility profiles of the 50 most accessible HERVL copies in diverse cell types from different brain regions at various developmental stages (data from Mannens et al., 2024^51^ and Trevino et al., 2021^59^), aligned to the HERVL consensus sequence. **e**, Schematic view of mouse embryo highlighting the cerebellum region at E13.5 (left) and sagittal section of the hindbrain along the plane shown on the left (right). Arrows indicate the migration paths of rhombic-lip-derived neuroblasts. **f**, *In situ* hybridization images from Allen Developing Mouse Brain Atlas^64^ (https://developingmouse.brain-map.org) showing expression of Atoh1 (left) and Nfib (right) in E13.5 mouse hindbrain. **g**, Jaccard similarity index matrix comparing the 30 most accessible HERVL copies between cell types highlighted in **a** and **d. h**, Examples of cCREs that are specific to cerebellar neuroblasts 1 (top), specific to hindbrain glutamatergic neuroblasts 4 (middle), or shared between these two cell types (bottom). Annotated HERVL copies are shown as gray bars. E, embryonic day.

The HERVL fragment at position 2,000-2,500 nt contains multiple NFI and E-box motifs (Fig. 2e), which are known to regulate neuronal differentiation across various regions of the nervous system^50,51,58^. This led us to investigate whether HERVL copies might also be co-opted during the development of other brain regions. We analyzed chromatin accessibility across multiple datasets, including a comprehensive atlas of the whole brain during the first trimester of human development^51^ and cerebral cortex datasets covering the second and third trimesters and postnatal development^59,60^ (Fig. 3c). Our analysis revealed that glutamatergic neuroblasts in the hindbrain (pons, cerebellum, and medulla), and not in other brain regions, show significant enrichment of accessible HERVL copies specifically at the 2,000-2,500 nt position (Fig. 3d). All these neuron lineages, including cerebellar granule cells and unipolar brush cells, are derived from the upper and lower rhombic lips based on their marker gene expression (Extended Data Fig. 4a-c), consistent with ATOH1 being a key regulator of these cell lineages^54,61–63^. Other cell types, including microglia and cortical radial glia, exhibit distinct accessibility patterns at different positions within the HERVL consensus sequence (Fig. 3d). Indeed, the HERVL 3,300-3,800 nt position, where microglia show enriched accessibility, contains two PU.1 binding motifs (GGAAGT) and displays high regulatory potential in NMF program 7 (microglia) (Extended Data Fig. 5a,b). To determine whether different cell types utilize the same or distinct HERVL copies, we computed a pairwise similarity matrix using the Jaccard index between the 30 most accessible HERVL copies across cell types. We identified three distinct clusters corresponding to microglia, cortical radial glia, and rhombic-lip-derived neuroblasts, as expected (Fig. 3e), indicating that cell types that co-opted different positions of the HERVL consensus sequence utilize largely non-overlapping sets of HERVL copies. Notably, we observed additional variability within rhombic-lip-derived neuroblasts (Fig. 3e,f), suggesting that these neuron lineages utilize partially distinct sets of HERVL copies depending on their developmental state and cell lineage identity, which could contribute to cell state-specific gene expression patterns associated with differentiation timing and different migratory pathways from the rhombic lips^54,61–63^. Collectively, these results demonstrate that the HERVL region originally encoding the polymerase protein of the endogenous retrovirus is preferentially co-opted for regulatory activity in rhombic-lip-derived neuroblasts, with different cell states within this lineage utilizing partially distinct sets of HERVL copies.

### HERVL-specific sequence substitutions create high regulatory potential in hindbrain neuroblasts

The 2,000-2,500 nt region of HERVL lies within the reverse transcriptase domain of the polymerase gene, one of the most conserved regions in otherwise fast-evolving endogenous retroviruses and frequently used to establish phylogenetic relationships among retroviruses^65,66^. The conserved nature of this region across related elements prompted us to investigate the sequence evolution of the HERVL subfamily to understand how this enhancer potential emerged in rhombic-lip-derived neuroblasts.

Comparative analysis of the consensus sequences of HERVL and seven TE subfamilies closely related to HERVL revealed that only HERVL exhibits significant chromatin accessibility enrichment in differentiating granule cells at the 2,000-2,500 nt position (Fig. 4a). Consistent with this observation, DeepCeREvo analysis for differentiating granule cells (NMF program 13) showed that other ERVL family TEs are assigned 36-58% lower prediction scores compared to HERVL (0.31-0.48 vs 0.74) at regions homologous to the HERVL 2,000-2,500 nt position (Fig. 4b). We identified multiple HERVL-specific substitutions in regions with high attribution scores that correspond to putative ATOH1 and NFI binding sites (Fig. 4c). The same patterns are also observed in individual genomic copies of these ERVL family transposons (Extended Data Fig. 6). Our findings show that the enhancer potential of HERVL in rhombic-lip-derived neuroblasts arises from unique sequence substitutions specific to this subfamily. To assess whether the high regulatory potential specific to HERVL translates to enhancer activity, we performed luciferase reporter assays in primary cultures of mouse granule cells isolated from P7 mouse cerebella and cultured *ex vivo* for 3 days. This experimental system has been validated for testing enhancer activity of both conserved and newly emerged CREs in differentiating granule cells^30^. We tested consensus sequences from 5 transposable elements closely related to HERVL, focusing on regions homologous to the HERVL 2,000-2,500 nt position. Among the tested sequences, the HERVL element exhibited significantly higher luciferase activity compared to fragments from dinucleotide-shuffled HERVL and other closely related ERVL family TEs in the reverse orientation, but not in the forward orientation (Fig. 4d; Supplementary Table 3). This suggests directional effects on its gene regulatory activity, aligning with earlier reports that enhancers are not fully orientation independent^67^. These results demonstrate that HERVL-specific sequence features confer functional enhancer activity that distinguishes it from other closely related elements, validating our computational predictions of its regulatory potential.

**Fig. 4:**
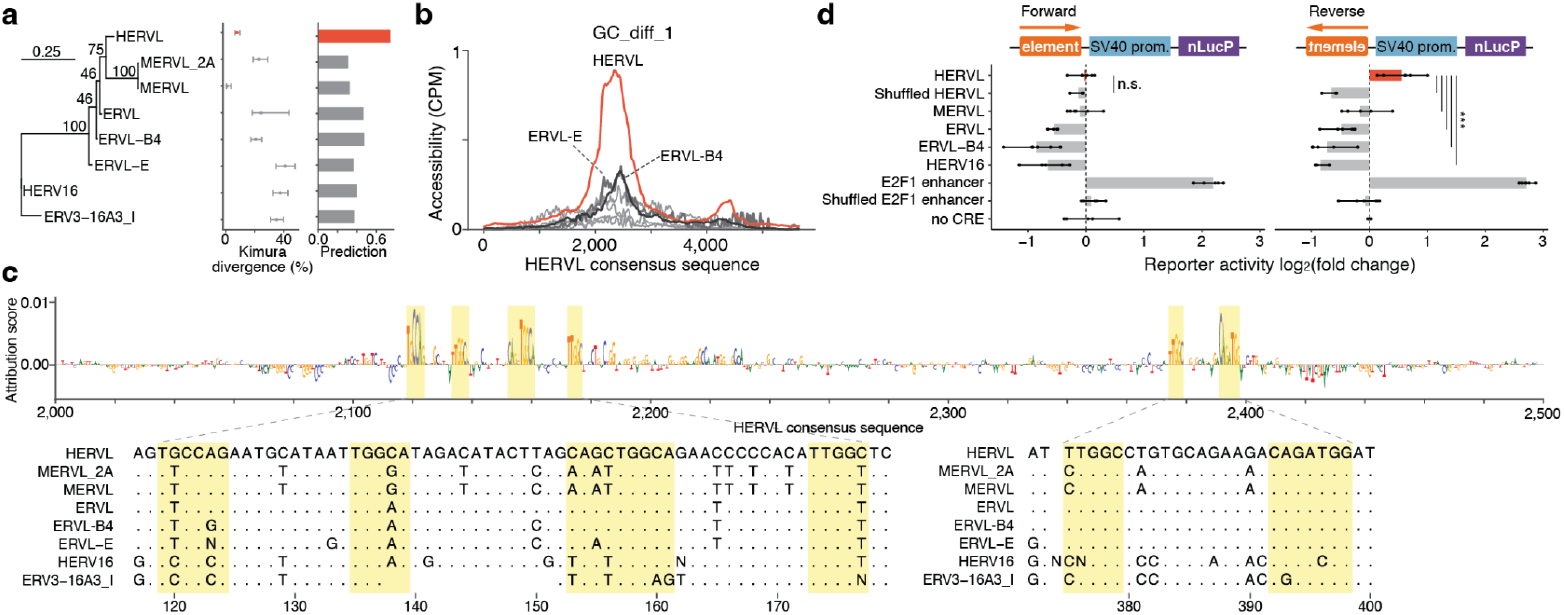
HERVL-specific sequence substitutions create unique enhancer activity. **a**, Phylogenetic relationship of HERVL-related ERVL family transposons (left) alongside their median divergence (with interquartile range) and DeepCeREvo prediction scores for the corresponding NMF program 13 (right), using regions orthologous to the 2,000-2,500 nt position in HERVL. Numbers in the internal nodes of the phylogenetic tree indicate bootstrap confidence values (%). **b**, Mean accessibility profile of the 50 most accessible copies of internal regions from ERVL family transposons related to HERVL. **c**, Multiple sequence alignment of consensus sequences of internal regions of ERVL family transposons compared to HERVL, highlighting regions with high DeepExplainer attribution scores. **d**, Luciferase reporter assays in mouse primary granule cells testing the enhancer activity of regions homologous to HERVL 2,000-2,500 nt position in other ERVL family transposons. The E2F1 fragment is a conserved enhancer active in differentiating granule cells^30^. Shuffled fragments were obtained by shuffling sequences while keeping the dinucleotide frequency. Candidate fragments were placed in front of the SV40 promoter in forward (left) and reverse (right) orientation. Bars and error bars display the mean normalized and scaled reporter activity and its range; points denote independent experiments. *P*-values against the HERVL fragment were estimated using linear models, corrected for multiple testing using the Benjamini-Hochberg method. ***, *P* < 0.001.

### Preservation of TF binding sequences and local chromatin context jointly determine HERVL accessibility

Despite having high regulatory potential, only a small subset of genomic HERVL copies (2.1%) are accessible in rhombic-lip-derived neuroblasts, raising the question of what distinguishes active from inactive elements. We found no significant differences between accessible and inaccessible copies in sequence divergence (a proxy for insertion age) or element length (Fig. 5a,b; Supplementary Table 4). Furthermore, phylogenetic analysis revealed that accessible and inaccessible copies distribute throughout the reconstructed evolutionary tree without forming distinct clusters, indicating their derivation from the same ancestral sequence (Fig. 5c; Extended Data Fig. 6). In contrast to these shared characteristics, DeepCeREvo prediction score analysis of the HERVL 2,000-2,500 position revealed significantly higher prediction scores for accessible copies compared to inaccessible ones (Fig. 5c,d; *P* < 10^−10^, Mann-Whitney *U* test; Supplementary Table 4), where the median score of accessible copies (0.72) was slightly lower than the consensus sequence (0.74). Consistently, sequences in high-attribution regions, corresponding to putative TF binding sites, show greater preservation in accessible copies than in inaccessible ones (Fig. 5e; Extended Data Fig. 7). These results indicate that local sequence features, particularly the preservation of TF binding sites present in the ancestral sequence, are the primary determinants of HERVL copy accessibility.

**Fig. 5:**
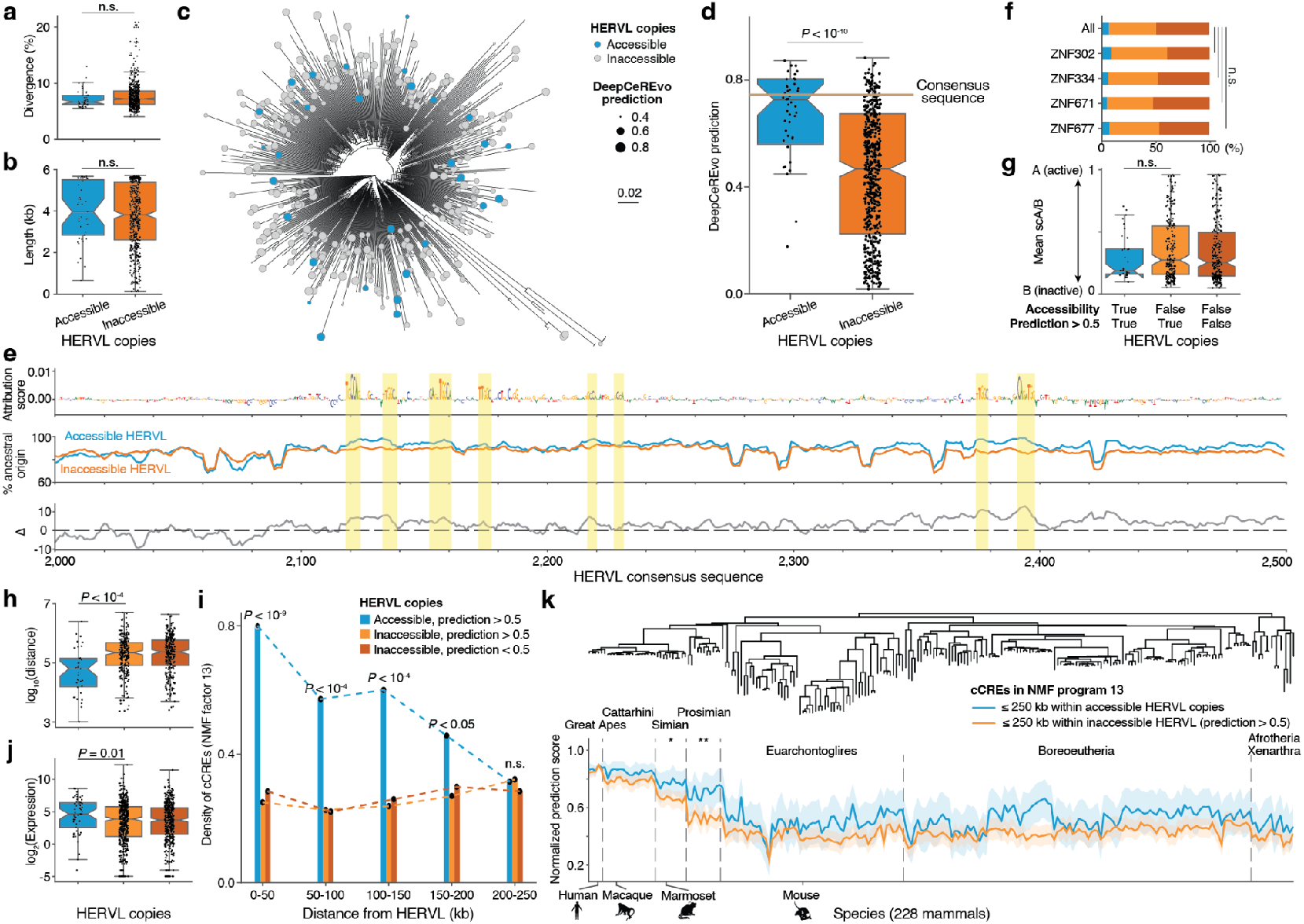
A subset of HERVL copies with high enhancer potential are accessible and involved in gene regulation in differentiating granule cells. **a, b**, Kimura divergence (**a**) and length (**b**) of accessible and inaccessible HERVL copies. **c**, Phylogenetic relationship of HERVL copies in the human genome, annotated with chromatin accessibility status in differentiating granule cells (GC_diff_1) and corresponding DeepCeREvo prediction scores (NMF program 13). **d**, Comparison of DeepCeREvo prediction scores (NMF program 13) between accessible and inaccessible HERVL copies in differentiating granule cells. **e**, Sequence conservation analysis of HERVL copies showing DeepExplainer attribution profile of the HERVL ancestral (consensus) sequence (top), percentage of nucleotides identical to the ancestral sequence in accessible versus inaccessible HERVL copies (middle), and differential conservation pattern between accessible and inaccessible copies (bottom). TF binding sites in the consensus sequence are highlighted. **f**, Percentage of HERVL copies within binding sites of the respective KZFPs. **g, h**, Compartment score (**g**) and log2 distance to the closest non-TE-derived cCREs specific to differentiating granule cells (**h**) of accessible and inaccessible HERVL copies. **i**, Density of non-TE-derived cCREs around HERVL copies. **j**, Expression levels of genes nearby accessible and inaccessible HERVL copies. **k**, DeepCeREvo prediction scores (NMF program 13) for human cCREs proximal to accessible and inaccessible HERVL copies, and their orthologous genomic regions across 227 placental mammalian species.

However, 47% of the inaccessible HERVL copies still received prediction scores greater than 0.5 (Fig. 5a,b), indicating that local sequence features alone cannot explain all cases of inaccessibility. We hypothesized that epigenetic silencing mechanisms mediated by KRAB domain-containing zinc finger proteins (KZFPs) or higher-order chromatin organization^68,69^ might be responsible for suppressing these high-potential elements. To test whether KZFP-mediated suppression explains inaccessibility, we analyzed binding sites of HERVL-targeting KZFPs overexpressed in HEK293T cells using previous datasets^70,71^ and showed that inaccessible HERVL copies are not significantly enriched for KZFP binding regions compared to accessible copies (Fig. 5f). We also examined higher-order chromatin effects by analyzing megabase-scale A/B compartments using single-cell 3D genome architecture data in developing human cerebellum^72^ and observed no significant differences in compartment positioning between accessible and inaccessible HERVL copies (Fig. 5g). These results suggest that neither KZFP-mediated suppression nor large-scale nuclear compartmentalization are primary determinants of HERVL accessibility for copies with high regulatory potential.

At a finer scale, however, the local chromatin landscape shows striking differences. Accessible HERVL copies are located significantly closer to non-TE-derived cell-type-specific cCREs (NMF program 13) than inaccessible ones, regardless of their predicted regulatory potential (*P* < 10^−4^, Mann-Whitney *U* test), with median distances of 64 kb for accessible and 221-234 kb for inaccessible copies (Fig. 5h). Consistently, these non-TE-derived cCREs are significantly enriched around accessible HERVL copies compared to inaccessible ones, with the strongest enrichment observed within 50 kb (Fisher’s exact *P* < 10^−9^) (Fig. 5i). Consistent with this pattern, genes proximal to accessible HERVL copies show higher expression levels than those near inaccessible copies (Fig. 5j; Supplementary Table 5). To determine if the local chromatin environment was already open prior to HERVL insertion, we examined the evolutionary histories of human cCREs around accessible HERVL copies, based on DeepCeREvo prediction scores in orthologous regions across 228 mammalian species^30^. This analysis revealed that orthologous regions of human cCREs (NMF program 13) nearby accessible HERVL copies have significantly higher prediction scores across primate lineages than those not close to accessible HERVL copies (Fig. 5k). This suggests that the chromatin regions surrounding accessible HERVL copies were likely already in an active state before HERVL insertion occurred. Collectively, these findings indicate that HERVL accessibility is determined by at least two complementary factors: insertion within pre-existing active chromatin neighborhoods containing other cell-type-specific regulatory elements and subsequent preservation of TF binding sites.

### Species-specific co-option of HERVL elements drives regulatory divergence in primate cerebellar development

HERVL elements underwent a major expansion in the primate lineage, with most copies inserted after the primate-rodent split^57^ (Fig. 6a). This evolutionary timing makes HERVL an excellent candidate for driving regulatory innovations specific to primate brain development. By analyzing our macaque and marmoset cerebellar datasets^30^, we find that HERVL copies are predominantly accessible in differentiating granule cells in both species, with 27 accessible copies identified in each (Extended Data Fig. 8a,b). The accessible copies in both species possess higher DeepCeREvo prediction scores compared to inaccessible copies (Extended Data Fig. 8c,d). These results indicate a conserved potential for HERVL co-option as CREs specific to rhombic-lip-derived neuroblasts across primates.

**Fig. 6:**
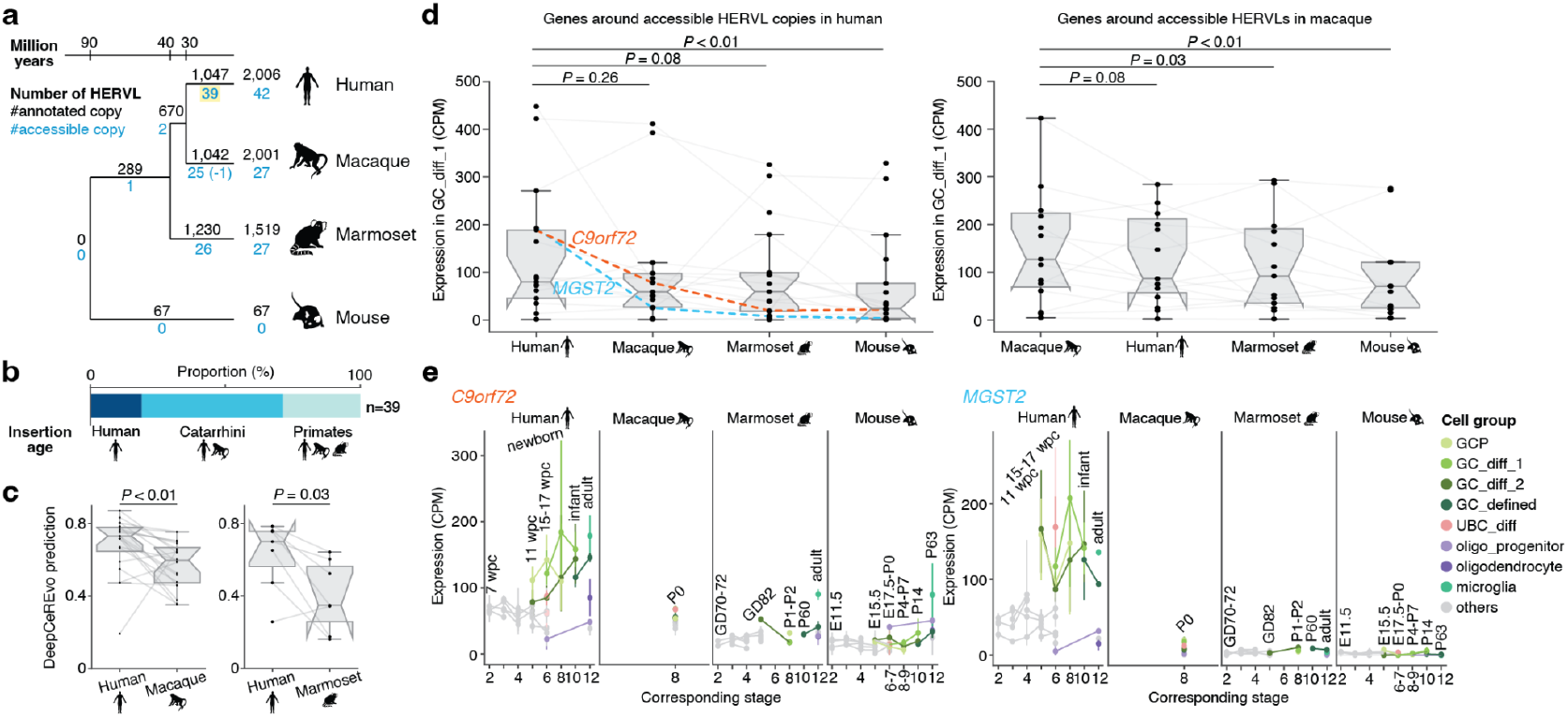
Species-specific accessible HERVL copies contributed to gene expression divergence during primate evolution. **a**, Number of annotated and accessible HERVL copies in differentiating granule cells (GC_diff_1) across human, macaque, marmoset, and mouse. **b**, Insertion age distribution of HERVL copies accessible in differentiating granule cells specifically in human. **c**, DeepCeREvo prediction scores for differentiating granule cells (NMF program 13) comparing human accessible HERVL copies with their corresponding orthologous sequences in macaque (left) and marmoset (right). **d**, Expression levels of 1:1 orthologous genes located near HERVL copies accessible in differentiating granule cells (GC_diff_1) in human (left) and macaque (right). **e**, Spatiotemporal expression profiles of *C9orf72* (left) and *MGST2* (right) across corresponding cell types and developmental stages in mammalian species. Lines represent mean expression and error bars indicate the range across biological replicates. CPM, counts per million. E, embryonic day; GD, gestational day; P, postnatal day; wpc, weeks post conception.

While the potential for HERVL co-option appears conserved across primates, the specific elements activated show remarkable lineage specificity. Comparative genomic analysis revealed that of 2,006 annotated HERVL copies in the human genome, 14% (n = 289) are shared across examined primates, 33% (n = 670) are restricted to catarrhines (i.e., apes and Old World Monkeys), and 52% (n = 1,047) are human-specific (Fig. 6a). Notably, the vast majority of accessible HERVL copies in each species are not orthologous to accessible copies in the other species (Fig. 6a). Among 39 HERVL copies specifically accessible in human, approximately 20% represent human-specific insertions, while the remainder have orthologous non-accessible HERVL copies in macaque or marmoset (Fig. 6b). The human-specific accessible HERVL copies have significantly higher DeepCeREvo prediction scores than the orthologous non-accessible HERVL copies in macaque and marmoset (*P* < 0.05, Wilcoxon signed-rank test) (Fig. 6c). Consistently, macaque- and marmoset-specific accessible HERVL copies also have higher prediction scores than the orthologous non-accessible human copies (Extended Data Fig. 8e,f), indicating species-specific accessibility of orthologous HERVL is determined by the species-specific preservation of ancestral TF binding sites. Furthermore, species-specific accessible HERVL copies are surrounded by non-TE-derived CREs specific to differentiating granule cells across all three primate species (Fig. 5h,i; Extended Data Fig. 8g,h). These results show that accessible HERVL copies are predominantly species-specific, occurring within distinct local chromatin environments and reflecting lineage-specific insertions and lineage-specific sequence divergence that affects ancestral TF binding sites.

To determine whether species-specific accessible HERVL copies influence gene regulation across primates, we examined expression patterns of nearby genes in differentiating granule cells. Genes associated with human-specific accessible HERVL copies frequently exhibit elevated expression compared to their orthologs in macaque and marmoset (Fig. 6d; Supplementary Table 5).

Several genes with particularly higher expression in human include *C9orf72*, whose dysfunction impairs neurodevelopment in human induced pluripotent stem cell (iPSC) and mouse models^73,74^, and *MGST2*, which produces a key signaling molecule in response to oxidative stress, to which cerebellar granule cells are particularly susceptible compared to other brain cell types^75^. The increased expression of these genes is specific to or most prominent in differentiating granule cells (GC_diff_1) (Fig. 6e), consistent with the high regulatory potential of HERVL in this cell type. Similar species-specific regulatory effects are observed in macaque, where genes near macaque-specific accessible HERVL copies also show higher expression compared to their 1:1 orthologous genes in other species (Fig. 6d). Collectively, these findings suggest that HERVL elements with intrinsic regulatory potential have been repeatedly and independently co-opted as species-specific enhancers during primate evolution, contributing to distinct chromatin accessibility landscapes and divergent gene expression patterns in differentiating granule cells in the cerebellum.

## DISCUSSION

In this study, we investigated the contributions of TEs to the chromatin accessibility landscape in cerebellum development and uncovered underlying evolutionary and genetic mechanisms using cross-species single-cell multi-omics atlases, along with a sequence-based deep-learning model predicting cell-type-specific chromatin accessibility. Notably, we identified that primate-specific endogenous retrovirus HERVL elements are preferentially co-opted as CREs specific to rhombic-lip-derived neuroblasts and drive gene expression evolution across the primate lineage.

Preferential co-option of TEs due to the presence of complex cis-regulatory sites in the ancestral forms of TEs has been widely documented, particularly in the LTR family during early embryogenesis^14,37,39^. LTR elements typically possess binding sites for TFs active during early development such as NANOG, SOX2, and OCT4, which drive the expression of their proviral sequences in the germline lineage, facilitating amplification of these elements within genomes through vertical transmission^38,47,52,53,76,77^. While these TF binding sites can be co-opted in stem cells or progenitors where similar TFs are active, the preferential co-option of TEs in differentiating or differentiated cell types has remained less characterized. This knowledge gap exists primarily because detailed molecular profiles such as histone modifications or TF binding patterns across diverse cellular contexts have been limited, constraining most analyses to the bulk tissue level^19,20,78^.

Our study addresses this limitation by combining single-cell chromatin accessibility profiles with a sequence-based deep-learning model that predicts cell-type-specific chromatin accessibility to identify seventeen TE fragments with high regulatory potential in specific cell types in developing cerebellum. Among these is HERVL, a primate-specific endogenous retrovirus that is co-opted in rhombic-lip-derived neuroblasts during hindbrain development. Notably, this co-option occurs through its polymerase gene region rather than the LTR sequences that typically serve as regulatory elements in retroviruses. Through detailed analysis of HERVL, we demonstrated that insertion events of these TEs can facilitate gene expression evolution, which could potentially contribute to the phenotypic evolution of cerebellum development, such as significant increases in neuron numbers, in the primate lineage. Furthermore, the framework developed here can be applied to any cellular context in any tissue at any developmental stage, potentially expanding our understanding of TE contributions to gene regulatory networks and to phenotypic evolution.

In addition to these preferential co-option events of specific TE subfamilies, our comprehensive analysis of single-cell chromatin accessibility data suggested that evolutionary constraints also have a broad impact on shaping the overall distribution of TE-derived cCREs across cell types. This conclusion is supported by consistent patterns observed in our study and others showing that TE-derived cCREs are enriched in less-constrained genomic contexts: distal elements rather than promoters, species-specific rather than conserved regions, cell-type-specific rather than pleiotropic sites^23^ and elements active in later rather than earlier developmental stages^51^. The reduced presence of TE-derived cCREs in early development might alternatively be explained by TE suppression mechanisms via KZFPs, whose expression is higher during prenatal development^79,80^. However, this suppression mechanism itself likely evolved in response to developmental constraints, as organisms would benefit from minimizing potentially deleterious effects of TE activity during more critical early developmental periods.

Our study has several limitations. First, our reliance on short-read sequencing makes it challenging to accurately profile chromatin accessibility in recently expanded long TEs with very low sequence divergence, such as L1HS, SINE-VNTR-Alu (SVA), and HERVK (HML-2)^81^. However, since these elements constitute only a fraction of annotated TEs, this limitation is unlikely to significantly affect our main conclusions. Additionally, our detailed characterization of preferential TE co-option is limited to HERVL, while the regulatory contributions of the other 16 identified TE fragments, and likely additional elements not captured by our conservative screening criteria, remain to be explored. Future systematic analysis of these elements would strengthen our understanding of TE-driven regulatory evolution.

Our work provides a new framework to study TE contributions to cis-regulatory landscapes and to gene regulatory evolution across different cell types, exemplified by HERVL co-option in hindbrain development across the primate lineage. This approach could extend beyond cerebellar development by integrating large-scale chromatin accessibility atlases with sequence-based deep-learning models^41,51,82,83^. Together, the framework and findings of this study may help inform future research into the contributions of TEs to the evolution of gene regulation and, ultimately, organismal phenotypes.

## ACKNOWLEDGEMENTS

We thank D. Odom, O. Stegle, R. Grand, P. Joshi, A. Stuart, A. Kondrashkina and all members of the Kaessmann lab for discussions and K. Hall, J. Schmidt and C. Schneider for assistance. The computational cluster bwForCluster of Heidelberg University Computational Center is supported by the state of Baden-Württemberg through bwHPC and the German Research Foundation (INST 35/1134-1 FUGG). The authors gratefully acknowledge the data storage service SDS@hd supported by the Ministry of Science, Research and the Arts Baden-Württemberg (MWK) and the German Research Foundation (DFG) through grant INST 35/1503-1 FUGG. T.Y was supported by the Takenaka Scholarship Foundation. I.S. was supported by an EMBO Scientific Exchange Grant (9231) and an EMBO Postdoctoral Fellowship (ALTF 769-2022). M.S. was supported by a Simons Foundation Autism Research Initiative (SFARI) Bridge to Independence Award (SFI-AN-AR-Independence Postdoctoral-00007139). This project has received funding from the European Research Council (ERC) under the European Union’s Horizon 2020 research and innovation programme (VerteBrain to H.K., grant agreement no. 101019268).

## Author contributions

T.Y., I.S., M.S., and H.K. conceived the study. T.Y. analyzed data with support from I.S. M.S. performed reporter assays. H.K. and I.S. supervised the study. H.K. provided funding. T.Y. drafted the manuscript, with critical review by I.S., M.S., and H.K. All authors approved its final version.

## Data availability

Previously published datasets are available at Array Express (E-MTAB-9765 and E-MTAB-10533)^28^ and the heiData repository (https://doi.org/10.11588/data/QDOC4E^29^ and https://heidata.uni-heidelberg.de/previewurl.xhtml?token=bea0c1aa-3110-4f3a-a98c-f10e65a407d0^30^).

## Code availability

All original code is available at https://gitlab.com/kaessmannlab/cerebellum_te.

## Competing interests

The authors declare no competing interests.

## METHODS

### Single-cell multi-omics atlas of cerebellum development

The snRNA-seq and snATAC-seq datasets across cerebellum development in human, marmoset, and mouse used in this study are the same as our previous study^30^. For cross-species comparison of cell-type-specific chromatin accessibilities, CREs specific to certain cell types in certain developmental stages in different species were grouped into programs based on non-negative matrix factorization (NMF), as described in our previous study^30^. In short, standardized pseudobulk accessibility matrices of human and mouse cis-regulatory elements (CREs) for 45 sample groups [CREs x samples] (combinations of cell types and developmental stages, which correspond in both human and mouse) were decomposed into two new matrices [CREs x programs] and [programs x samples] via a predetermined number of programs. The number of programs was determined to be 18 based on the reconstruction error and the distance between CREs from human and mouse in the same group. We assigned human and mouse highly variable CREs to each program based on the [CREs x programs] matrix (CRE loadings). For marmoset, the CRE loading matrix ([CREs x programs]) was obtained by using non-negative least squares on the [CREs x samples] matrix in marmoset and the [programs x samples] matrix obtained in the human and mouse analysis. Similarly, all highly accessible peaks in human, marmoset, and mouse were assigned to NMF programs based on the same transformation and the same CRE loading threshold.

### Overlap between TEs and CREs in mammalian cerebellum development

To identify the overlap between transposable elements and the CREs accessible in cerebellum development, we intersected CREs with annotated transposable elements in human, marmoset, and mouse genomes. We downloaded non-overlapping repeat annotations for human (hg38), macaque (rheMac8), marmoset (ASM275486v1), and mouse (mm10) from the Dfam database (v3.8)^31^. Repeat annotations for ASM275486v1 marmoset assembly were transferred to the calJac4 assembly using UCSC liftOver with a custom chain file generated by nf-LO^84^. Only repeat annotations whose classes belong to LTR, LINE, SINE, and DNA were retained. Since the LTR class contains both long terminal repeats and internal elements of endogenous retroviruses (ERVs), and these sequences show significant differences, this class was further classified into LTR and ERV-int subclasses. This classification was mostly based on the description in the Dfam database when available, which was the case in nearly 95% of the transposons in the LTR class. Otherwise, it was based on a length threshold of 2,000 bp. We intersected these curated transposable element annotations with CRE annotations using bedtools (v2.30.0)^85^ and only retained overlaps longer than 250 bp (i.e., half of the cCRE length).

### Enrichment of TEs in CREs

Enrichment of TEs in CREs was computed as described in the previous study^16^. Specifically, we calculated the enrichment by dividing the fraction of overlaps between TE annotations and CREs of interest (relative to the total number of CREs of interest) by the fraction of genomic bases covered by the TE annotations of interest relative to the total genome size. For visualization purposes, we assigned a log2 enrichment value of −10 to TE subfamilies with no overlap with cCREs, which is lower than the enrichment value of any subfamily that does exhibit cCRE overlap.

### Inferring the Age of Transposon Subfamilies

We inferred the age of transposon subfamilies based on the divergence of their respective copies in the genome. Since each copy in the same subfamily accumulates mutations after its insertion, the divergence from its original form (inferred through consensus sequence reconstruction) increases over time. We computed the median Kimura divergence score of transposon copies from their consensus sequences and used this as a measure of transposon subfamily age.

### Assessing regulatory potential of TE subfamilies

We assessed the regulatory potential of TE subfamilies using the deep-learning model DeepCeREvo, which predicts chromatin accessibility in specific cell states (NMF programs) in cerebellum development by detecting combinations of cell-type-specific TF binding sites^30^. We obtained consensus sequences of all TE subfamilies in human and mouse using famdb.py (v1.0.2) on the FamDB HDF5 database (partition 0) from Dfam database (v3.8). We only considered TE subfamilies with at least 10 overlaps with highly variable cCREs in at least one cell group, resulting in 127 subfamilies. Since DeepCeREvo requires 500-bp input sequences, we created sliding windows of 500 bp with 100-bp steps to generate TE fragments for assessment.

Given that TE-derived cCREs are enriched in cell-type-specific elements, we developed an approach to compute the regulatory potential specific to one or a few cell groups. First, we transformed the DeepCeREvo outputs by applying the inverse sigmoid function (logit) and then the softmax function to convert values into a probability distribution across cell groups. This transformation enhances cell=type-specificity by generating high values for elements specific to certain cell types while assigning low values to elements with ubiquitously high accessibility. Next, we masked CpG dinucleotides from input sequences. This step was necessary because consensus TE fragments typically contain excess CpG dinucleotides compared to non-CpG-island genomic sequences, which is one of sequence features of promoters^86^. Finally, to account for sub-optimal proto-motif sequences in TE fragments, we computed regulatory potential by averaging transformed prediction scores from the original sequence and 25 sequences which had single nucleotide variants introduced to the original sequence and showed the highest regulatory potential. Specifically, we generated all possible single nucleotide variants of the original sequence, evaluated their regulatory potential, and selected the top 25 highest-scoring variants to include in our averaging calculation.

To determine statistical significance, we shuffled each TE fragment 1,000 times and computed regulatory potential for these shuffled sequences. By comparing regulatory potential between original and shuffled fragments, we calculated empirical P values and applied the Benjamini-Hochberg procedure for multiple-test correction. Recognizing that tests between adjacent TE fragments and between closely related TE subfamilies are not independent, we used the number of TE fragments divided by 10 as the effective number of tests to establish conservative thresholds controlling the false discovery rate (FDR) at 5%.

### Computing positional enrichment of TE contributions to cCREs

We evaluated how much sequences derived from ancestral TE copies contribute to cell-type-specific cCREs. This analysis complements our regulatory potential screening, as we would expect TE fragments with high regulatory potential to contribute more significantly to cCREs of their corresponding cell types. To quantify this contribution, we first intersected highly variable cCREs with TE annotations using bedtools intersect (v2.30.0)^85^. We then mapped these intersections to corresponding consensus sequences using MAFFT (v7.505) with the --addfragments option^87^. For each TE fragment, we computed the number of bases aligned to cCREs and counted it as an overlap if ≥ 100 bases were aligned. To establish background expectations, we applied the same procedure to a chromatin accessibility atlas containing cell types from the whole human body at both fetal and adult stages^41^ Finally, we calculated positional enrichment of each TE fragment by dividing the proportion of highly variable cCREs in cerebellum development overlapping each TE fragment by the proportion of background cCREs overlapping the same fragment.

### Mapping chromatin accessibility across HERVL consensus and identifying accessible copies in differentiating granule cells

To analyze the chromatin accessibility patterns of HERVL elements, we first constructed a multiple sequence alignment (MSA) of all 2,006 annotated HERVL copies in the human genome using MAFFT (v7.505) with the --addfragments option^87^, aligning them to the HERVL consensus sequence as reference. For each HERVL copy, we mapped chromatin accessibility data from various cell types to its corresponding position in the MSA using pyBigWig (v0.3.18) on accessibility tracks in bigWig format. This approach enabled us to generate consensus-based accessibility profiles for HERVL elements across different cell types. To identify HERVL copies with accessible chromatin specifically at the 2,000-2,500 nt position in differentiating granule cells, we computed the total accessibility score for each HERVL copy at this region. Based on the distribution of these scores, we established 0.3 as the threshold value to classify HERVL elements as accessible at this critical position, allowing us to distinguish between accessible and inaccessible HERVL copies in this cell type.

### HERVL accessibility across developmental stages and brain regions

To investigate HERVL chromatin accessibility patterns throughout brain development and across different neuroanatomical regions, we integrated data from multiple sources. We obtained chromatin accessibility tracks for cell types in the first trimester of human brain development from the CATlas database^41,51^. Recognizing that different brain regions develop at varying rates, with hindbrain structures typically developing earlier than other regions, we incorporated additional chromatin accessibility datasets spanning second trimester to adult stages of cerebral cortex development^59,60^. Since the accessibility data from Herring et al. was originally mapped to the hg19 assembly, we converted these files to hg38 coordinates using CrossMap (v0.7.0)^88^ to ensure consistency across all analyses. For each cell type represented in these datasets, we used the corresponding chromatin accessibility tracks in bigWig format to compute relative accessibility profiles of HERVL copies based on their alignment positions within the consensus sequence in the same way as the previous section.

### Constructing phylogenetic tree of ERVL family transposons

We first built a multiple sequence alignment of the consensus sequences of the ERV-int region from 12 ERVL family transposons from human and MERVL and MERVL-2A from mouse using MAFFT (v7.505)^87^. We identified sequences orthologous to the HERVL 2,000-2,500 nt position and retained subfamilies with overlap longer than 250 bp. With these orthologous sequences, we again built a multiple sequence alignment. We removed poorly aligned regions from the sequence alignment using trimAl (v1.4) with a -gappyout parameter^89^ and built a phylogenetic tree using IQ-TREE2 (v2.3.4) with the default parameters^90^. We excluded 5 ERVL family transposons (HERVL74, ERVL40, HERVL66, HERVL32, HERVL1, and ERVL47) due to large divergence from HERVL and reran the analysis to obtain the final phylogenetic tree of ERVL family transposons at the HERVL 2,000-2,500 nt position. We used treeio (v1.18.1) to load tree objects and ggtree (v3.2.1) to visualize the inferred phylogenetic tree^91,92^.

### Constructing phylogenetic tree of ERVL family transposon copies in the human genome

To construct phylogenetic tree of ERVL family transposon copies, we obtained genomic coordinates of ERVL family transposons (HERVL, ERVL, ERVL-B4, ERVL-E, HERV16, and ERV3-16A3-I) in the human genome, which were identified as closely related to HERVL based on our consensus sequence analysis. We retained only annotated TE copies longer than 2,000 bp to ensure analysis of structurally intact and well-annotated elements, minimizing the inclusion of heavily fragmented or degenerate sequences. We then built multiple sequence alignments and constructed the phylogenetic tree using the same methodology as described for consensus sequences, with one additional criterion: we only included copies that overlapped with the 2,000-2,500 nt position of the HERVL consensus by at least 400 bp to ensure the reliability of the reconstructed phylogenetic tree. For our HERVL-specific phylogenetic analysis, we applied more lenient filtering criteria, including annotated copies longer than 250 bp that overlapped with the HERVL 2,000-2,500 nt position by at least 250 bp.

### Luciferase reporter assays

Reporter assays were performed and analysed essentially as described previously^30^. Briefly, primary granule cell cultures were prepared from P7 mouse cerebella and cultured for 3 days *in vitro* (DIV3) in the presence of 200 nM Smoothened Agonist to support proliferation of granule cell progenitors. ERVL family transposon sequences were synthesized based on consensus sequences from the Dfam database, with ambiguous nucleotides (‘N’) resolved by alignment of all copies in the human genome to the consensus sequence and selection of the most frequently observed nucleotides at each position. Sequences were cloned into pNL1.2[NlucP]-based vectors upstream of an SV40 promoter in both forward and reverse orientations. Cells were transfected at DIV2 using FuGENE HD with 95 ng of test constructs and 30 ng of firefly luciferase normalizer plasmid. Luciferase activities were measured 28-30 hours post-transfection using the Nano-Glo Dual-Luciferase Reporter Assay System. For the estimation of enhancer activities, only wells with firefly luciferase signals at least 3-fold above background were included. Statistical analysis was performed using linear mixed models with R packages lme4 (v1.1-36)^93^, lmerTest (v3.1-3)^94^, and pbkrtest (v.0.5-0.1)^95^, with element, orientation, and their interaction as fixed variables and independent experiments as random variables. The data did not support inclusion of a random intercept for experiment (singular fit), resulting in a model equivalent to a fixed-effects linear model. *P*-values were corrected for multiple comparisons using the Benjamini-Hochberg method. All sequences and values are provided in Supplementary Table 3.

### KZFP binding site analysis

We used a public database on KZFP binding sites from chromatin immunoprecipitation and lambda exonuclease digestion (ChIP-exo) experiments on HEK293T cell lines overexpressing different KZFPs with a hemagglutinin (HA) tag^70,71^. We obtained a list of KZFPs, (ZNF157, ZNF254, ZNF302, ZNF485, ZNF671, ZNF707, ZNF677, ZNF793, ZNF334, and PRDM9), whose binding sites are enriched in HERVL (padj.binomial < 0.05). KZFPs that are barely expressed (CPM < 10) in differentiating granule cells (GC_diff_1) (ZNF157, ZNF485, ZNF707, and PRDM9) were excluded. ChIP-exo peaks for each KZFP were intersected with the coordinates of HERVL copies that had the 2,000-2,500 position of the consensus sequence. KZFPs with less than 5 binding sites with these HERVL copies (ZNF254 and ZNF793) were also excluded from the following analysis. We then counted the number of HERVL copies with different features that have binding sites for each KZFP and computed the fraction.

### Single cell chromosomal compartment analysis

We analyzed the chromatin compartmentalization of HERVL elements in human cerebellar development using single-cell chromatin A/B compartment (scA/B) datasets obtained from^72^. A/B compartments represent regions of open euchromatin and closed heterochromatin, respectively. Based on the comparative analysis of cell type abundance profiles across developmental stages, we identified granule cell structural stage S1 as corresponding most closely to the transcriptomic cell type GC_diff_1, with both populations showing peak enrichment during the newborn stage. To characterize the chromatin environment of HERVL elements, we first calculated the genome-wide mean scA/B scores across all 707 cells annotated to the granule cell S1 population. We then determined the midpoints of annotated HERVL copies. The chromatin compartmentalization status of each HERVL element was assessed by extracting the scA/B score at its corresponding midpoint location.

### Local chromatin environment analysis

To investigate how the local chromatin environment influences HERVL copy accessibility in differentiating granule cells, we analyzed the spatial relationship between HERVL elements and cCREs. We focused on cCREs specifically accessible in NMF program 13 (differentiating granule cells) that did not overlap with any annotated TEs ≥ 250 bp in length. For each HERVL copy, we determined the distance to the nearest non-TE-derived, NMF program 13-specific cCRE using bedtools closest (v2.30.0) with the pybedtools wrapper (v0.9.0)^85,96^.

To characterize the broader regulatory landscape, we calculated the density of these non-TE-derived cCREs around HERVL copies using bedtools window (v2.30.0) with a window size of 250 kb^85,96^. We divided this window into 50 kb bins and counted the number of cCREs in each bin. The density was calculated by dividing these counts by the number of HERVL copies in each class, providing a normalized measure of regulatory element distribution around different categories of HERVL elements.

### Analysis of gene expression in proximity to HERVL elements

To examine the potential regulatory influence of HERVL elements on nearby genes, we assessed the relationship between HERVL accessibility and the expression of genes within a 500 kb window (±250 kb) centered on each HERVL copy. We extended the center position of each HERVL element using bedtools slop (v2.30.0) and identified genes with transcription start sites (TSS) within this window through bedtools intersect (v2.30.0). For each identified gene, we computed the mean expression level across all samples classified as differentiating granule cells (GC_diff_1), enabling us to evaluate whether accessible HERVL elements were associated with differential expression of neighboring genes.

### Identification of orthologous HERVL copies between species

To investigate the evolutionary impact of HERVL insertions on chromatin accessibility and gene expression in differentiating granule cells across primate lineages, we identified orthologous HERVL elements between species. Using the UCSC liftOver tool with default parameters, we mapped human HERVL genomic coordinates to their corresponding 1:1 orthologous regions in macaque, marmoset, and mouse genomes. We then employed bedtools intersect (v2.30.0) to determine which of these syntenic regions contained annotated HERVL elements in the target species^85^. To robustly define orthology, we only retained orthologous elements that were explicitly annotated as HERVL in the non-human genome and possessed a length at least 20% of their human counterparts, thus filtering out severely truncated or degraded elements.

### Evaluation of species-specific HERVL contributions to cross-species gene expression divergence

To investigate whether species-specific accessible HERVL elements contribute to evolutionary divergence of gene expression patterns, we analyzed a set of 1:1:1:1 orthologous protein-coding genes conserved across human, macaque, marmoset, and mouse genomes. We quantile-normalized the mean expression values of these genes in differentiating granule cells (GC_diff_1) across all four species to facilitate direct cross-species comparisons. To identify genes potentially influenced by human-specific HERVL insertions, we focused on the most highly expressed gene within a 500 kb window (±250 kb) surrounding each HERVL copy, reducing technical noise while prioritizing genes more likely under regulatory influence of these elements.

## Extended Data Figures

**Extended Data Fig. 1:**
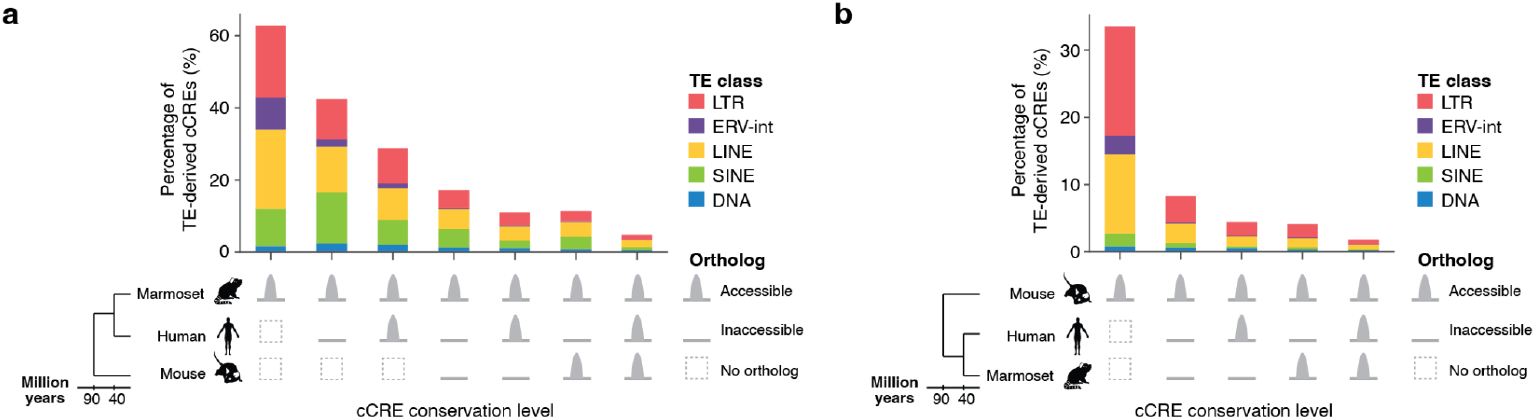
Transposable element contributions to cCREs during marmoset and mouse cerebellar development. **a, b**, Percentage of TE-derived cCREs during marmoset (**a**) and mouse (**b**) cerebellar development across sequence conservation levels between human, marmoset, and mouse.

**Extended Data Fig. 2:**
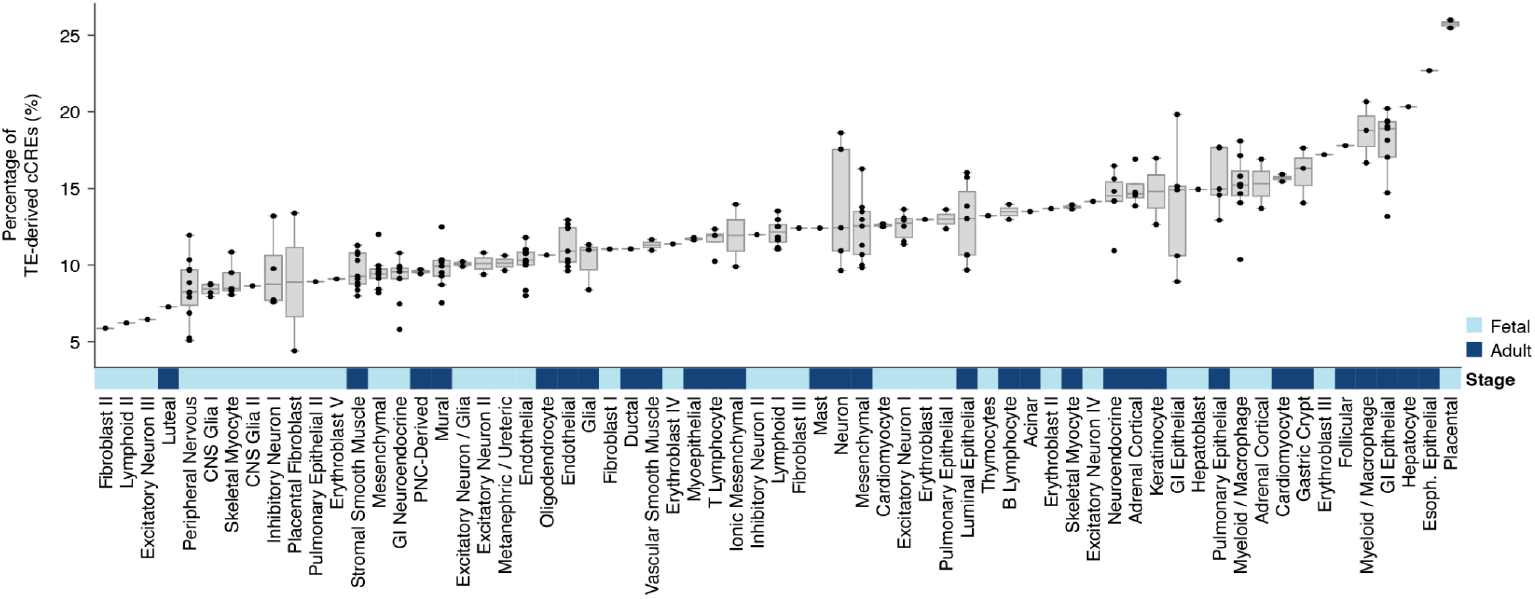
Fraction of TE-derived cCREs varies across fetal and adult human cell types. Percentage of TE-derived cCREs across individual cell types (subclusters, dots) grouped by major cell type clusters. Samples are separated into fetal and adult developmental stages. Data from Zhang et al., 2021^41^ reanalyzed.

**Extended Data Fig. 3:**
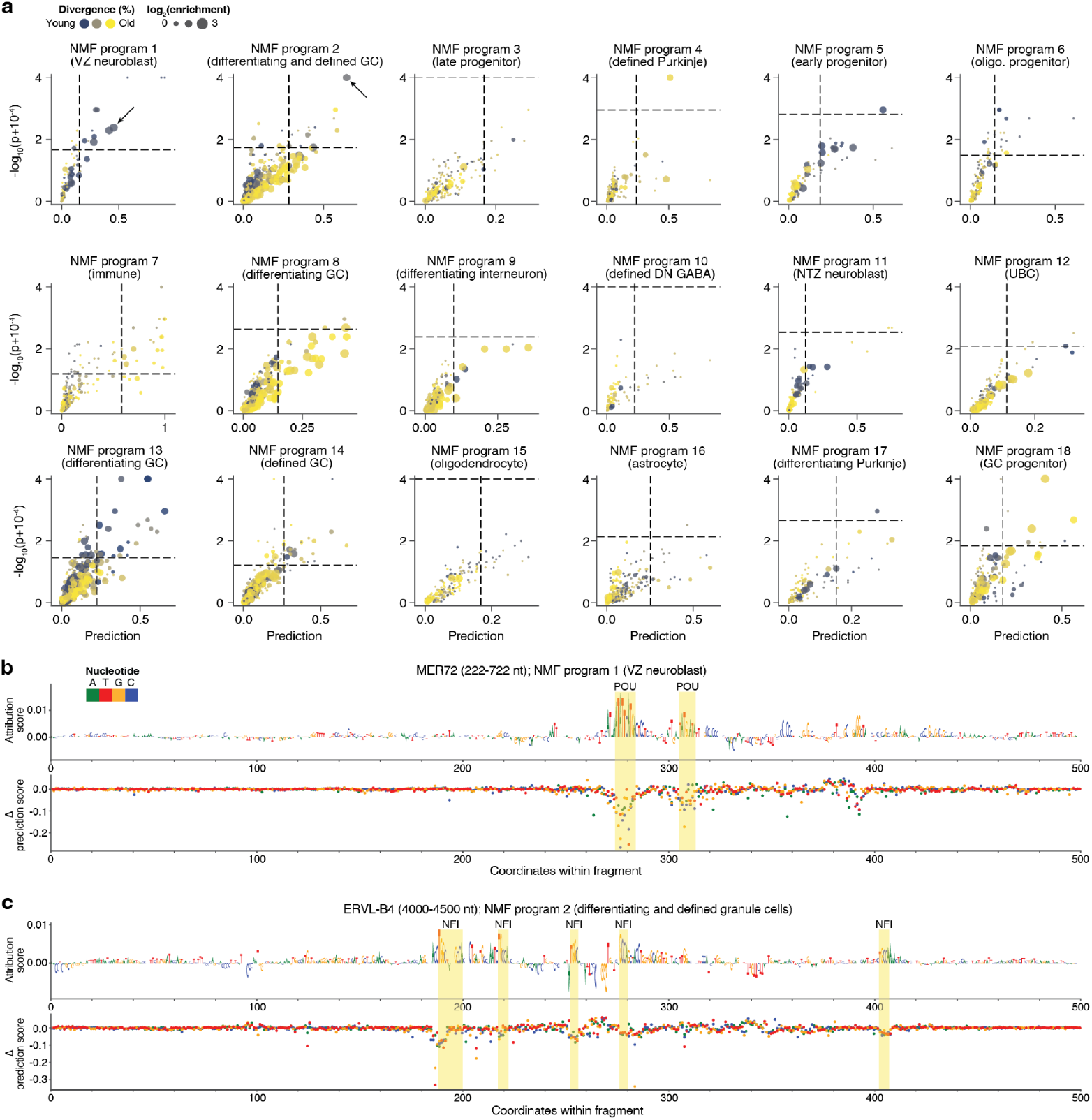
*In silico* screening of transposable elements with high regulatory potential in human cerebellar development. **a**, Screening of TE fragments based on regulatory potentials and associated *P*-values, alongside enrichment in cell-type-specific cCREs for each NMF program. The regulatory potential threshold was set at the top 5th percentile. The *P*-value threshold was determined using the Benjamini–Hochberg procedure to control the false discovery rate (FDR) at 5%, adjusted for the effective number of tests. **b, c**, DeepExplainer attribution and *in silico* mutagenesis profiles for top candidates in NMF program 1 (VZ neuroblasts) (**b**) and NMF program 2 (differentiating granule cells) (**c**), which are indicated by arrows in **a**. VZ, ventricular zone; NTZ, nuclear transitory zone; GC, granule cell; UBC, unipolar brush cell; DN GABA, deep nuclei GABAergic neuron.

**Extended Data Fig. 4:**
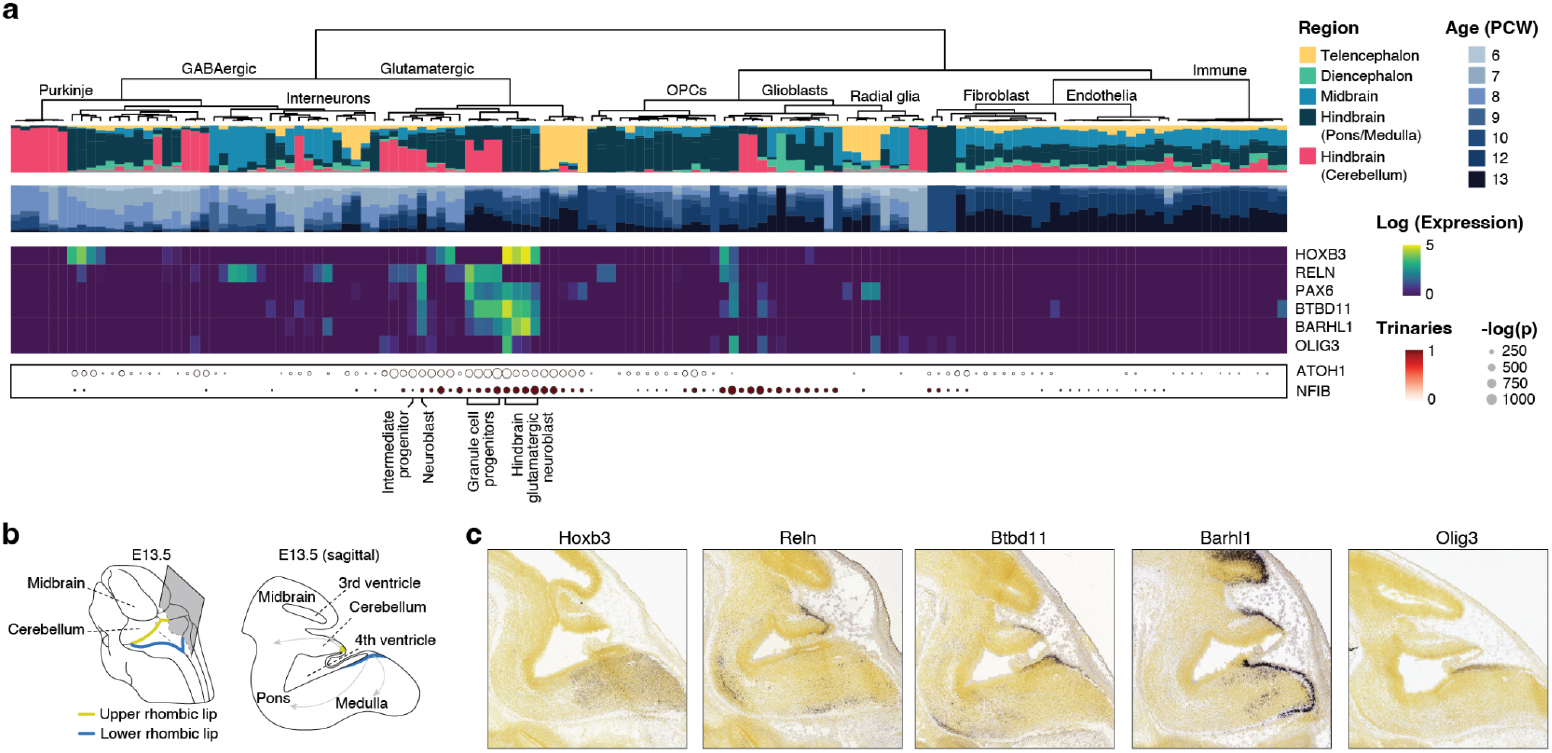
Anatomical characterization of cell types with accessible HERVL elements during first trimester human brain development. **a**, Top to bottom: hierarchical clustering dendrogram showing relationships between cell types based on highly variable cCREs; anatomical region distribution showing the proportion of each cell type across brain regions; developmental age distribution showing temporal profiles of each cell type; expression heatmap of selected marker genes across cell types; TF binding motif enrichment analysis with dot size representing statistical significance (*P* values) and color intensity representing expression levels of the corresponding TFs (trinarization score represents a probabilistic measure of gene expression ranging from 0-1). Highlighted cell types show HERVL accessibility (Fig. 3d). Data from Mannens et al., 2024^51^ reanalyzed. **b**, Schematic view of mouse embryo highlighting the cerebellum region at E13.5 (left) and sagittal section of the hindbrain along the plane shown on the left (right). Arrows indicate the migration paths of rhombic-lip-derived neuroblasts. **c**, *In situ* hybridization images from Allen Developing Mouse Brain Atlas^64^ (https://developingmouse.brain-map.org) showing expression of marker genes in E13.5 mouse hindbrain.

**Extended Data Fig. 5:**
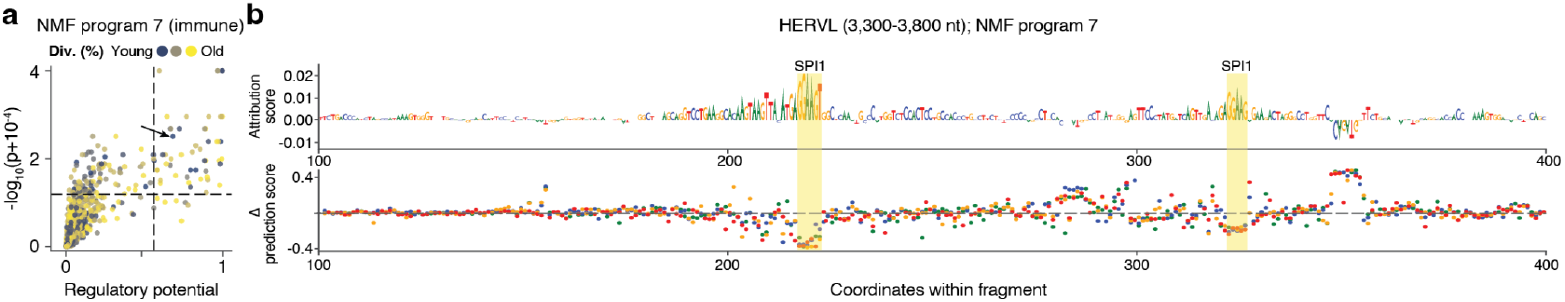
Characterization of HERVL at 3,300-3,800 nt position in microglia. **a**, Regulatory potentials and associated *P*-values in NMF program 7 (microglia) for all TE fragments. The regulatory potential threshold was set at the top 5th percentile. The *P*-value threshold was determined using the Benjamini–Hochberg procedure to control the false discovery rate (FDR) at 5%, adjusted for the effective number of tests. **b**, DeepExplainer attribution and *in silico* mutagenesis profiles in NMF program 7 (microglia) for HERVL at 3,300-3,800 nt position, which is indicated by an arrow in **a**.

**Extended Data Fig. 6:**
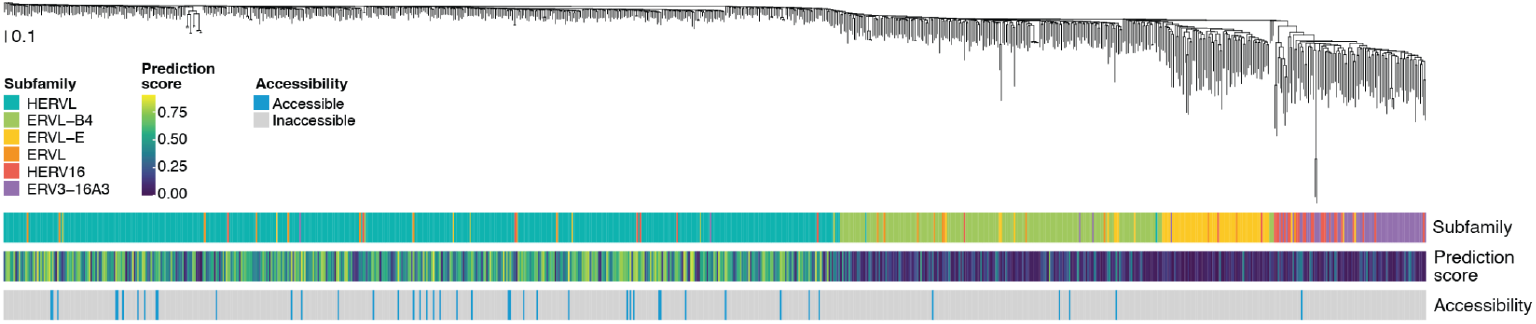
Phylogenetic relationships of ERVL family transposons in the human genome. Phylogenetic tree of 852 ERVL family transposons in the human genome at regions orthologous to the HERVL 2,000-2,500 nt position. For each copy, annotated subfamily, DeepCeREvo prediction score for NMF program 13 (differentiating granule cells), and accessibility in GC_diff_1 are shown below.

**Extended Data Fig. 7:**
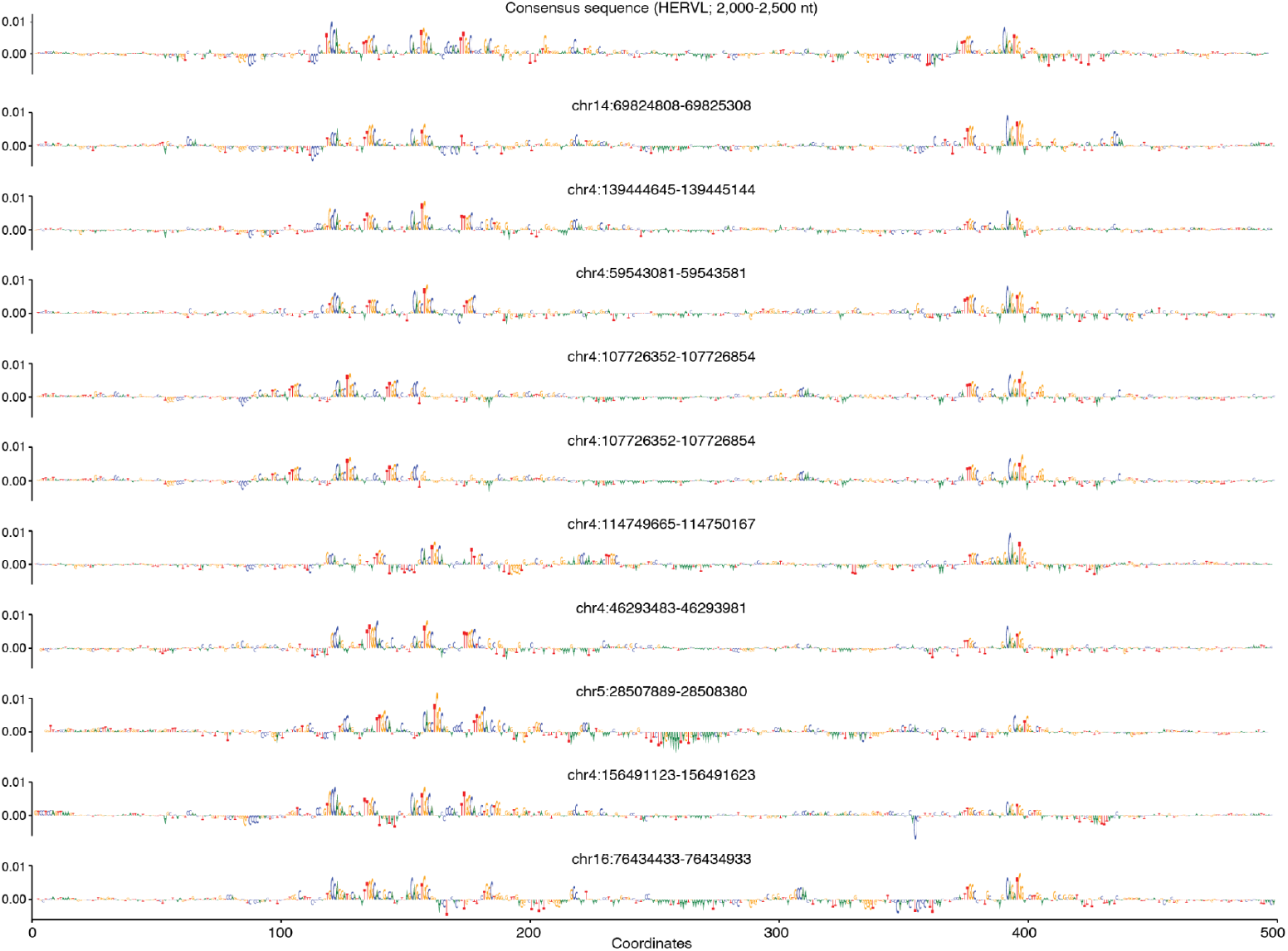
Accessible HERVL copies in the human genome. DeepExplainer attribution profiles of 10 HERVL copies accessible in GC_diff_1 with the highest DeepCeREvo prediction scores.

**Extended Data Fig. 8:**
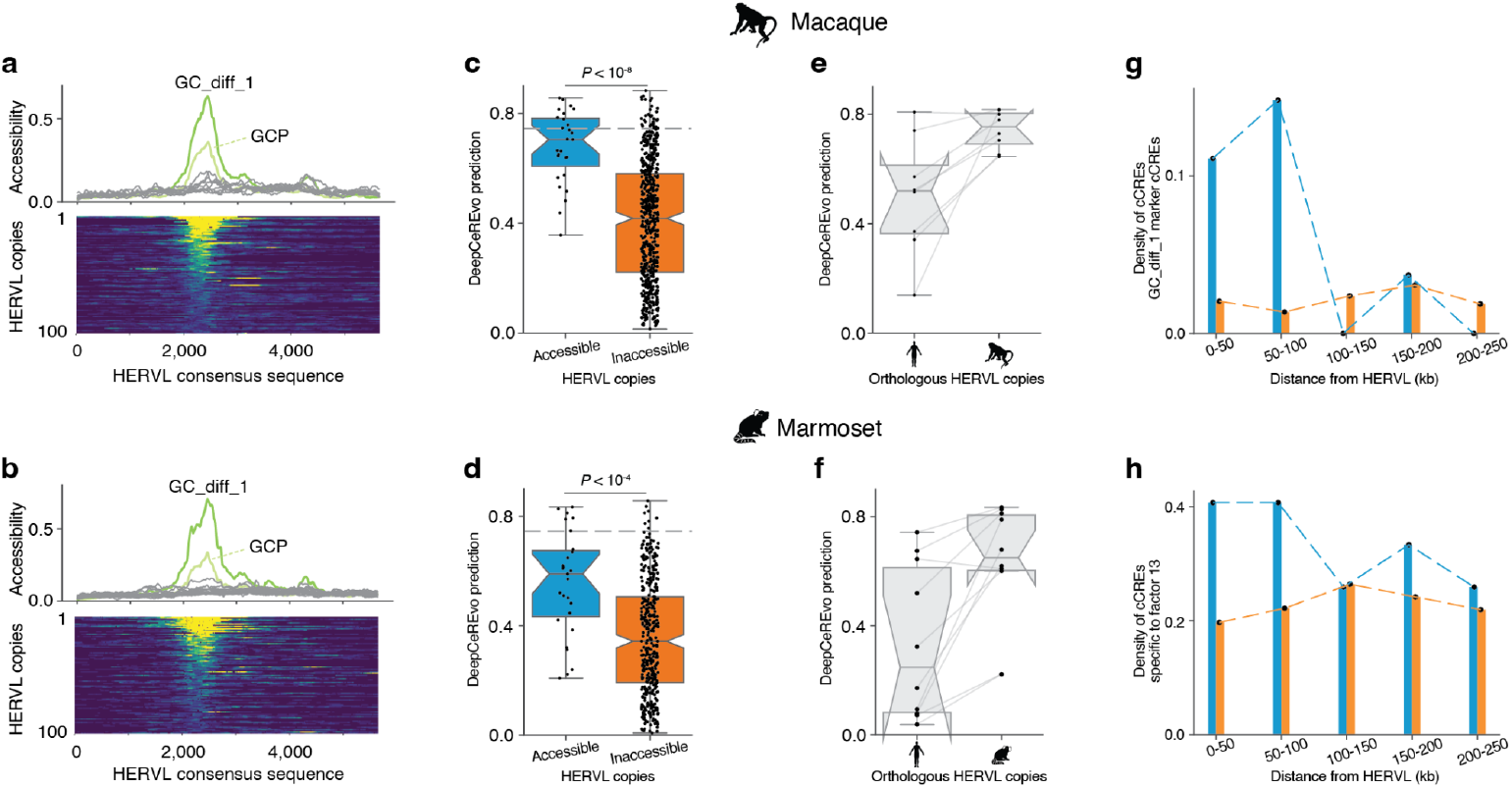
Analyses on HERVL copies in macaque and marmoset. **a, b**, Mean chromatin accessibility profiles of the 50 most accessible HERVL copies across cell groups in cerebellum development (left) and Chromatin accessibility patterns of the 100 most accessible HERVL copies in the 2,000-2,500 nt position in differentiating granule cells (GC_diff_1), aligned to the consensus sequence (right) in macaque (**a**) and marmoset (**b**). **c, d**, DeepCeREvo prediction scores between accessible and inaccessible HERVL copies in macaque (**c**) and marmoset (**d**). **e, f**, Density of non-TE-derived cCREs specific to differentiating granule cells around accessible and inaccessible HERVL copies in macaque (**e**) and marmoset (**f**). **g, h**, Expression levels of genes nearby accessible and inaccessible HERVL copies in macaque (**g**) and marmoset (**h**).

## Supplementary Information

**Supplementary Table 1**. Enrichment of TE-derived cCREs across NMF programs in human cerebellum development.

**Supplementary Table 2**. Regulatory potential screening results for 500 bp TE fragments using DeepCeREvo.

**Supplementary Table 3**. Reporter construct sequences and luciferase assay results for selected TE fragments.

**Supplementary Table 4**. Genomic features and accessibility status of HERVL copies in the human genome.

**Supplementary Table 5**. Expression levels of genes proximal to accessible and inaccessible HERVL copies.

## Notes

https://gitlab.com/kaessmannlab/cerebellum_te

### Competing Interest Statement

The authors have declared no competing interest.

## REFERENCES

1. Herculano-Houzel, S. Coordinated scaling of cortical and cerebellar numbers of neurons. Front. Neuroanat. 4, 12 (2010).

2. Herculano-Houzel, S. The human brain in numbers: a linearly scaled-up primate brain. Front. Hum. Neurosci. 3, 31 (2009).

3. King, M. C. & Wilson, A. C. Evolution at two levels in humans and chimpanzees. Science 188, 107–116 (1975).

4. Carroll, S. B. Evo-devo and an expanding evolutionary synthesis: a genetic theory of morphological evolution. Cell 134, 25–36 (2008).

5. Wilson, M. D. et al. Species-specific transcription in mice carrying human chromosome 21. Science 322, 434–438 (2008).

6. Shlyueva, D., Stampfel, G. & Stark, A. Transcriptional enhancers: from properties to genome-wide predictions. Nat. Rev. Genet. 15, 272–286 (2014).

7. Bourque, G. Transposable elements in gene regulation and in the evolution of vertebrate genomes. Curr. Opin. Genet. Dev. 19, 607–612 (2009).

8. Rebollo, R., Romanish, M. T. & Mager, D. L. Transposable elements: an abundant and natural source of regulatory sequences for host genes. Annu. Rev. Genet. 46, 21–42 (2012).

9. de Souza, F. S. J., Franchini, L. F. & Rubinstein, M. Exaptation of transposable elements into novel cis-regulatory elements: is the evidence always strong? Mol. Biol. Evol. 30, 1239–1251 (2013).

10. McClintock, B. Controlling elements and the gene. Cold Spring Harb. Symp. Quant. Biol. 21, 197–216 (1956).

11. Britten, R. J. & Davidson, E. H. Gene regulation for higher cells: a theory. Science 165, 349–357 (1969).

12. Davidson, E. H. & Britten, R. J. Regulation of gene expression: Possible role of repetitive sequences. Science 204, 1052–1059 (1979).

13. Sundaram, V. & Wang, T. Transposable element mediated innovation in gene regulatory landscapes of cells: Re-visiting the ‘gene-battery’ model. Bioessays 40, (2018).

14. Sundaram, V. et al. Widespread contribution of transposable elements to the innovation of gene regulatory networks. Genome Res. 24, 1963–1976 (2014).

15. Pehrsson, E. C., Choudhary, M. N. K., Sundaram, V. & Wang, T. The epigenomic landscape of transposable elements across normal human development and anatomy. Nat. Commun. 10, 5640 (2019).

16. Du, A. Y., Chobirko, J. D., Zhuo, X., Feschotte, C. & Wang, T. Regulatory transposable elements in the encyclopedia of DNA elements. Nat. Commun. 15, 7594 (2024).

17. Lynch, V. J., Leclerc, R. D., May, G. & Wagner, G. P. Transposon-mediated rewiring of gene regulatory networks contributed to the evolution of pregnancy in mammals. Nat. Genet. 43, 1154–1159 (2011).

18. Sundaram, V. et al. Functional cis-regulatory modules encoded by mouse-specific endogenous retrovirus. Nat. Commun. 8, 14550 (2017).

19. Chuong, E. B., Elde, N. C. & Feschotte, C. Regulatory evolution of innate immunity through co-option of endogenous retroviruses. Science 351, 1083–1087 (2016).

20. Judd, J., Sanderson, H. & Feschotte, C. Evolution of mouse circadian enhancers from transposable elements. Genome Biol. 22, 193 (2021).

21. Smit, A. F. Interspersed repeats and other mementos of transposable elements in mammalian genomes. Curr. Opin. Genet. Dev. 9, 657–663 (1999).

22. de Koning, A. P. J., Gu, W., Castoe, T. A., Batzer, M. A. & Pollock, D. D. Repetitive elements may comprise over two-thirds of the human genome. PLoS Genet. 7, e1002384 (2011).

23. Simonti, C. N., Pavlicev, M. & Capra, J. A. Transposable element exaptation into regulatory regions is rare, influenced by evolutionary age, and subject to pleiotropic constraints. Mol. Biol. Evol. 34, 2856–2869 (2017).

24. Medstrand, P., van de Lagemaat, L. N. & Mager, D. L. Retroelement distributions in the human genome: Variations associated with age and proximity to genes. Genome Res. 12, 1483–1495 (2002).

25. Jurka, J., Kohany, O., Pavlicek, A., Kapitonov, V. V. & Jurka, M. V. Duplication, coclustering, and selection of human Alu retrotransposons. Proc. Natl. Acad. Sci. U. S. A. 101, 1268–1272 (2004).

26. Platt, R. N., 2nd, Vandewege, M. W. & Ray, D. Mammalian transposable elements and their impacts on genome evolution. Chromosome Res. 26, 25–43 (2018).

27. Nellåker, C. et al. The genomic landscape shaped by selection on transposable elements across 18 mouse strains. Genome Biol. 13, R45 (2012).

28. Sarropoulos, I. et al. Developmental and evolutionary dynamics of cis-regulatory elements in mouse cerebellar cells. Science 373, (2021).

29. Sepp, M. et al. Cellular development and evolution of the mammalian cerebellum. Nature 625, 788–796 (2024).

30. Sarropoulos, I. et al. The evolution of gene regulation in mammalian cerebellum development. bioRxiv 2025.03.14.643248 (2025) doi:10.1101/2025.03.14.643248.

31. Storer, J., Hubley, R., Rosen, J., Wheeler, T. J. & Smit, A. F. The Dfam community resource of transposable element families, sequence models, and genome annotations. Mob. DNA 12, 2 (2021).

32. Lynch, M. & Walsh, B. The Origins of Genome Architecture. (Oxford University Press, New York, NY, 2007).

33. Andrews, G. et al. Mammalian evolution of human cis-regulatory elements and transcription factor binding sites. Science 380, eabn7930 (2023).

34. Zemke, N. R. et al. Conserved and divergent gene regulatory programs of the mammalian neocortex. Nature 624, 390–402 (2023).

35. Wicker, T. et al. A unified classification system for eukaryotic transposable elements. Nat. Rev. Genet. 8, 973–982 (2007).

36. Cohen, C. J., Lock, W. M. & Mager, D. L. Endogenous retroviral LTRs as promoters for human genes: a critical assessment. Gene 448, 105–114 (2009).

37. Jacques, P.-É., Jeyakani, J. & Bourque, G. The majority of primate-specific regulatory sequences are derived from transposable elements. PLoS Genet. 9, e1003504 (2013).

38. Thompson, P. J., Macfarlan, T. S. & Lorincz, M. C. Long terminal repeats: From parasitic elements to building blocks of the transcriptional regulatory repertoire. Mol. Cell 62, 766–776 (2016).

39. Ito, J. et al. Systematic identification and characterization of regulatory elements derived from human endogenous retroviruses. PLoS Genet. 13, e1006883 (2017).

40. Hyacinthe, J. & Bourque, G. Transposable elements impact the regulatory landscape through cell type specific epigenomic associations. bioRxiv (2024) doi:10.1101/2024.08.07.606967.

41. Zhang, K. et al. A single-cell atlas of chromatin accessibility in the human genome. Cell 184, 5985–6001.e19 (2021).

42. Senft, A. D. & Macfarlan, T. S. Transposable elements shape the evolution of mammalian development. Nat. Rev. Genet. 22, 691–711 (2021).

43. Frost, J. M. et al. Regulation of human trophoblast gene expression by endogenous retroviruses. Nat. Struct. Mol. Biol. 30, 527–538 (2023).

44. Cardoso-Moreira, M. et al. Gene expression across mammalian organ development. Nature 571, 505–509 (2019).

45. Wells, J. N. & Feschotte, C. A Field Guide to Eukaryotic Transposable Elements. Annu. Rev. Genet. 54, 539–561 (2020).

46. Chuong, E. B., Elde, N. C. & Feschotte, C. Regulatory activities of transposable elements: from conflicts to benefits. Nat. Rev. Genet. 18, 71–86 (2017).

47. Fueyo, R., Judd, J., Feschotte, C. & Wysocka, J. Roles of transposable elements in the regulation of mammalian transcription. Nat. Rev. Mol. Cell Biol. 23, 481–497 (2022).

48. Johnson, W. E. Origins and evolutionary consequences of ancient endogenous retroviruses. Nat. Rev. Microbiol. 17, 355–370 (2019).

49. Notwell, J. H., Chung, T., Heavner, W. & Bejerano, G. A family of transposable elements co-opted into developmental enhancers in the mouse neocortex. Nat. Commun. 6, 6644 (2015).

50. Sagner, A. et al. A shared transcriptional code orchestrates temporal patterning of the central nervous system. PLoS Biol. 19, e3001450 (2021).

51. Mannens, C. C. A. et al. Chromatin accessibility during human first-trimester neurodevelopment. Nature 1–8 (2024).

52. Rodriguez-Terrones, D. & Torres-Padilla, M.-E. Nimble and ready to mingle: Transposon outbursts of early development. Trends Genet. 34, 806–820 (2018).

53. Sundaram, V. & Wysocka, J. Transposable elements as a potent source of diverse cis-regulatory sequences in mammalian genomes. Philos. Trans. R. Soc. Lond. B Biol. Sci. 375, 20190347 (2020).

54. Ben-Arie, N. et al. Math1 is essential for genesis of cerebellar granule neurons. Nature 390, 169–172 (1997).

55. Flora, A., Klisch, T. J., Schuster, G. & Zoghbi, H. Y. Deletion of Atoh1 disrupts Sonic Hedgehog signaling in the developing cerebellum and prevents medulloblastoma. Science 326, 1424–1427 (2009).

56. Klisch, T. J. et al. In vivo Atoh1 targetome reveals how a proneural transcription factor regulates cerebellar development. Proc. Natl. Acad. Sci. U. S. A. 108, 3288–3293 (2011).

57. Bénit, L., Lallemand, J. B., Casella, J. F., Philippe, H. & Heidmann, T. ERV-L elements: a family of endogenous retrovirus-like elements active throughout the evolution of mammals. J. Virol. 73, 3301–3308 (1999).

58. Harris, L., Genovesi, L. A., Gronostajski, R. M., Wainwright, B. J. & Piper, M. Nuclear factor one transcription factors: Divergent functions in developmental versus adult stem cell populations. Dev. Dyn. 244, 227–238 (2015).

59. Trevino, A. E. et al. Chromatin and gene-regulatory dynamics of the developing human cerebral cortex at single-cell resolution. Cell 184, 5053–5069.e23 (2021).

60. Herring, C. A. et al. Human prefrontal cortex gene regulatory dynamics from gestation to adulthood at single-cell resolution. Cell 185, 4428–4447.e28 (2022).

61. Wang, V. Y., Rose, M. F. & Zoghbi, H. Y. Math1 expression redefines the rhombic lip derivatives and reveals novel lineages within the brainstem and cerebellum. Neuron 48, 31–43 (2005).

62. Machold, R. & Fishell, G. Math1 is expressed in temporally discrete pools of cerebellar rhombic-lip neural progenitors. Neuron 48, 17–24 (2005).

63. Butts, J. C. et al. A single-cell transcriptomic map of the developing Atoh1 lineage identifies neural fate decisions and neuronal diversity in the hindbrain. Dev. Cell 59, 2171–2188.e7 (2024).

64. Thompson, C. L. et al. A high-resolution spatiotemporal atlas of gene expression of the developing mouse brain. Neuron 83, 309–323 (2014).

65. Boeke, J. D. & Stoye, J. P. Retrotransposons, endogenous retroviruses, and the evolution of retroelements. in Retroviruses (Cold Spring Harbor Laboratory Press, Cold Spring Harbor (NY), 1997).

66. Vargiu, L. et al. Classification and characterization of human endogenous retroviruses; mosaic forms are common. Retrovirology 13, 7 (2016).

67. Georgakopoulos-Soares, I. et al. Transcription factor binding site orientation and order are major drivers of gene regulatory activity. Nat. Commun. 14, 2333 (2023).

68. Dixon, J. R., Gorkin, D. U. & Ren, B. Chromatin Domains: The Unit of Chromosome Organization. Mol. Cell 62, 668–680 (2016).

69. Robson, M. I., Ringel, A. R. & Mundlos, S. Regulatory Landscaping: How Enhancer-Promoter Communication Is Sculpted in 3D. Mol. Cell 74, 1110–1122 (2019).

70. Imbeault, M., Helleboid, P.-Y. & Trono, D. KRAB zinc-finger proteins contribute to the evolution of gene regulatory networks. Nature 543, 550–554 (2017).

71. de Tribolet-Hardy, J. et al. Genetic features and genomic targets of human KRAB-zinc finger proteins. Genome Res. 33, 1409–1423 (2023).

72. Tan, L. et al. Lifelong restructuring of 3D genome architecture in cerebellar granule cells. Science 381, 1112–1119 (2023).

73. Settembre, C. et al. Systemic inflammation and neurodegeneration in a mouse model of multiple sulfatase deficiency. Proc. Natl. Acad. Sci. U. S. A. 104, 4506–4511 (2007).

74. Hendricks, E. et al. The C9ORF72 repeat expansion alters neurodevelopment. Cell Rep. 42, 112983 (2023).

75. Harrison, J. F. et al. Oxidative stress-induced apoptosis in neurons correlates with mitochondrial DNA base excision repair pathway imbalance. Nucleic Acids Res. 33, 4660–4671 (2005).

76. Kunarso, G. et al. Transposable elements have rewired the core regulatory network of human embryonic stem cells. Nat. Genet. 42, 631–634 (2010).

77. Barakat, T. S. et al. Functional dissection of the enhancer repertoire in human embryonic stem cells. Cell Stem Cell 23, 276–288.e8 (2018).

78. Nishihara, H. Retrotransposons spread potential cis-regulatory elements during mammary gland evolution. Nucleic Acids Res. 47, 11551–11562 (2019).

79. Rowe, H. M. & Trono, D. Dynamic control of endogenous retroviruses during development. Virology 411, 273–287 (2011).

80. Playfoot, C. J. et al. Transposable elements and their KZFP controllers are drivers of transcriptional innovation in the developing human brain. Genome Res. 31, 1531–1545 (2021).

81. Lanciano, S. & Cristofari, G. Measuring and interpreting transposable element expression. Nat. Rev. Genet. 21, 721–736 (2020).

82. Li, Y. E. et al. A comparative atlas of single-cell chromatin accessibility in the human brain. Science 382, eadf7044 (2023).

83. Liu, B. B. et al. Dissecting regulatory syntax in human development with scalable multiomics and deep learning. bioRxiv 2025.04.30.651381 (2025) doi:10.1101/2025.04.30.651381.

84. Talenti, A. & Prendergast, J. nf-LO: A Scalable, Containerized Workflow for Genome-to-Genome Lift Over. Genome Biol. Evol. 13, (2021).

85. Quinlan, A. R. & Hall, I. M. BEDTools: a flexible suite of utilities for comparing genomic features. Bioinformatics 26, 841–842 (2010).

86. Deaton, A. M. & Bird, A. CpG islands and the regulation of transcription. Genes Dev. 25, 1010–1022 (2011).

87. Katoh, K., Misawa, K., Kuma, K.-I. & Miyata, T. MAFFT: a novel method for rapid multiple sequence alignment based on fast Fourier transform. Nucleic Acids Res. 30, 3059–3066 (2002).

88. Zhao, H. et al. CrossMap: a versatile tool for coordinate conversion between genome assemblies. Bioinformatics 30, 1006–1007 (2014).

89. Capella-Gutiérrez, S., Silla-Martínez, J. M. & Gabaldón, T. trimAl: a tool for automated alignment trimming in large-scale phylogenetic analyses. Bioinformatics 25, 1972–1973 (2009).

90. Minh, B. Q. et al. IQ-TREE 2: New Models and Efficient Methods for Phylogenetic Inference in the Genomic Era. Mol. Biol. Evol. 37, 1530–1534 (2020).

91. Wang, L.-G. et al. Treeio: An R package for phylogenetic tree input and output with richly annotated and associated data. Mol. Biol. Evol. 37, 599–603 (2020).

92. Yu, G., Smith, D. K., Zhu, H., Guan, Y. & Lam, T. T.-Y. Ggtree: An r package for visualization and annotation of phylogenetic trees with their covariates and other associated data. Methods Ecol. Evol. 8, 28–36 (2017).

93. Bates, D., Mächler, M., Bolker, B. & Walker, S. Fitting linear mixed-effects models Usinglme4. J. Stat. Softw. 67, (2015).

94. Kuznetsova, A., Brockhoff, P. B. & Christensen, R. H. B. LmerTest package: Tests in linear mixed effects models. J. Stat. Softw. 82, (2017).

95. Halekoh, U. & Højsgaard, S. A Kenward-Roger approximation and parametric bootstrap methods for tests in linear mixed models - TheRPackagepbkrtest. J. Stat. Softw. 59, (2014).

96. Dale, R. K., Pedersen, B. S. & Quinlan, A. R. Pybedtools: a flexible Python library for manipulating genomic datasets and annotations. Bioinformatics 27, 3423–3424 (2011).

